# Temporal and spatial variability of mudflat and mangrove foraminiferal eDNA assemblages and its implication for sea-level reconstruction

**DOI:** 10.1101/2025.08.28.672876

**Authors:** Zhaojia Liu, Nicole S. Khan, Howard K.Y. Yu, Magali Schweizer, Jennifer S. Walker, Celia Schunter

## Abstract

Reconstructing relative sea level (RSL) is essential for understanding coastal evolution and mitigating impacts of climate change. Foraminiferal assemblages are established proxies for past sea levels, but their composition can vary seasonally and spatially, affecting the reliability of morphological reconstructions. Environmental DNA (eDNA) enables high-resolution, non-invasive monitoring of foraminiferal communities and supports high-precision RSL reconstruction. However, the spatiotemporal stability of eDNA assemblages in (sub)tropical intertidal zones—and their influence on RSL reconstruction—remains uncertain. We conducted a two-year eDNA monitoring study at three intertidal stations of varying tidal elevation in Hong Kong, sampling during both dry and wet seasons to assess variability in mangrove and mudflat environments. Mid-mangrove eDNA communities exhibited temporal and spatial stability. In contrast, eDNA assemblages in mudflat and upper-mangrove environments, particularly among monothalamous foraminifera taxa, showed pronounced seasonal shifts primarily driven by environmental changes. Despite this variability in the upper-mangrove, eDNA-based elevation estimates in mangrove consistently aligned with observed elevations (within 95% credible intervals), demonstrating reliability of RSL reconstructions in these environments. However, samples from mudflats, especially during the wet season, exhibited an overprediction bias, reflecting their heightened sensitivity to seasonal and exogenous eDNA inputs. These findings highlight the need to account for seasonal and environmental variability in eDNA-based RSL reconstruction. Stable mangroves are optimal for transfer functions, while transitional/mudflat zones require caution due to higher variability. Our study provides guidance for foraminiferal eDNA application in complex, dynamic coastal settings.

## 1. Introduction

Coastal ecosystems face unprecedented pressures from climate change, requiring robust indicators to monitor environmental change and guide management decisions. Foraminifera are valuable ecological indicators in coastal environments because they are sensitive to environmental stressors, including salinity, pollution, temperature, nutrient availability, eutrophication, and climate-driven changes (Bouchet et al., 2021; Frontalini et al., 2009; Koukousioura et al., 2011; Romano et al., 2021; Scott et al., 2007; Youssef et al., 2021). They also contribute to essential ecosystem services through nutrient cycling and sediment processes (Langlet et al., 2020; Piña-Ochoa et al., 2010; Risgaard-Petersen et al., 2006), making their monitoring valuable for integrated coastal management assessing both current ecosystem health and historical environmental change. An important application of foraminiferal indicators is to reconstruct past relative sea level (RSL) changes, which is essential for understanding coastal evolution and predicting future changes (Khan et al., 2017; Tan et al., 2023; Xiong et al., 2018). Foraminifera are sensitive to the frequency and duration of tidal inundation, forming vertically-zoned assemblages along elevation gradients in intertidal habitats (Berkeley et al., 2009a; Scott & Medioli, 1978). Traditional RSL reconstructions rely on morphological identification of foraminiferal tests in sediment cores, calibrated with modern training sets that relate community composition to tidal elevation (Barnett et al., 2016; Rush et al., 2021).

A key challenge is the variability in surface foraminiferal assemblages, which fluctuate both temporally (seasonal and interannual) and spatially at sub-meter scales (Berkeley et al., 2008; Buragohain & Ghosh, 2021; Buzas, 1970; Buzas et al., 2015; Horton & Edwards, 2003; Kemp et al., 2011; Richirt et al., 2020). These variability can bias RSL reconstructions, especially because modern training sets are often based on single samples collected at one time point per site (Edwards & Horton, 2000; Horton & Culver, 2008; Horton & Edwards, 2006; Kemp et al., 2013; Scott & Medioli, 1978; Yu et al., 2025). Living foraminiferal populations, typically identified by rose Bengal staining (Murray & Bowser, 2000; Walton, 1952), are particularly sensitive to these fluctuations, while dead assemblages are more stable over time and space (Hayward et al., 1996; Kemp et al., 2011; Murray & Alve, 2000; Scott et al., 2007; Walker et al., 2020). Consequently, RSL studies use dead assemblages for calibration (Hawkes et al., 2010; Horton, 1999; Yu et al., 2025), and recent work shows that accounting for temporal and spatial variability can improve RSL reconstruction accuracy (Walker et al., 2020).

Environmental DNA (eDNA) analysis has recently emerged as a promising alternative to conventional morphological approaches for studying foraminifera (Pawlowski et al., 2014). The application of eDNA techniques to foraminiferal community analysis enhances the ability to detect subtle ecological shifts in the transitional ecosystems before visible habitat alterations occur (Singer et al., 2023). Furthermore, foraminiferal eDNA shows distinct vertical zonation correlated with tidal elevation, enabling the use of transfer functions for late Holocene RSL reconstruction (Liu et al., 2025). eDNA can detect taxa that do not readily fossilize and are often missed in traditional studies, providing a more comprehensive and high-resolution view of foraminiferal diversity (Lejzerowicz et al., 2013a; Pawłowska et al., 2014; Singer et al., 2023) and potentially improving the predictive power of RSL reconstructions (Liu et al., 2025).

Despite these advantages, eDNA methods may also face similar challenges to morphological approaches. eDNA assemblages may not only comprise extracellular DNA (Nagler et al., 2022) — originating locally and dominated by DNA from deceased local populations (autochthonous) or transported from elsewhere (allochthonous sources) (Ellegaard et al., 2020; Nagler et al., 2022) — but also propagules (Brinkmann et al., 2023; Fouet et al., 2024; Singer et al., 2023), and living individual (Angeles et al., 2020) sources, all of which are subject to seasonal variation. eDNA is especially sensitive to subtle and small-scale variations in community composition (Bista et al., 2017; Matsuoka et al., 2021; Singer et al., 2023; Urabe et al., 2025), and has been widely applied in temporal biomonitoring (Brinkmann et al., 2023; He et al., 2019; Pawlowski et al., 2016) and biodiversity assessment (Bakker et al., 2019; Gu et al., 2023; Lejzerowicz et al., 2014). Thus, foraminiferal eDNA assemblages are likely to exhibit pronounced seasonal and small-scale spatial variability. However, the extent and implications of this variability for sea-level reconstructions remain poorly understood and have not been systematically assessed.

To address this knowledge gap, we investigated the temporal and spatial variability of foraminiferal eDNA assemblages and their effects on transfer function-based RSL reconstructions in mangrove and mudflat environments in Hong Kong. Over two years, we conducted semi-annual monitoring to: (1) assess seasonal dynamics and spatial variation in eDNA assemblages across tidal elevations; (2) identify environmental drivers of temporal variation; and (3) determine how seasonality influences the accuracy of eDNA-based RSL reconstructions. Our findings reveal clear seasonality in foraminiferal eDNA assemblages in coastal wetlands and provide recommendations for improving eDNA sampling strategies for RSL reconstruction.

## 2. Material and methods

### 2.1 Study area

Our study was conducted in the Mai Po Nature Reserve, situated on the eastern fringe of the Pearl River Delta (PRD), South China (Fig. 1B). The PRD is one of the world’s largest and most dynamic river deltas, supporting high biodiversity within its extensive networks of rivers, estuaries, and intertidal wetlands (WWF, 2020). Rapid urbanization and industrial development, particularly in neighboring cities Shenzhen and Hong Kong, have influenced regional hydrology, sediment dynamics, and water quality throughout Deep Bay and the Mai Po wetlands (Sun et al., 2017). The region experiences semidiurnal tides with a great diurnal amplitude of 1.96 m (from mean lower low water [MLLW] to mean higher high water [MHHW]), recorded by the nearest tide gauge station at Tsim Bei Tsui.

**Figure 1.**
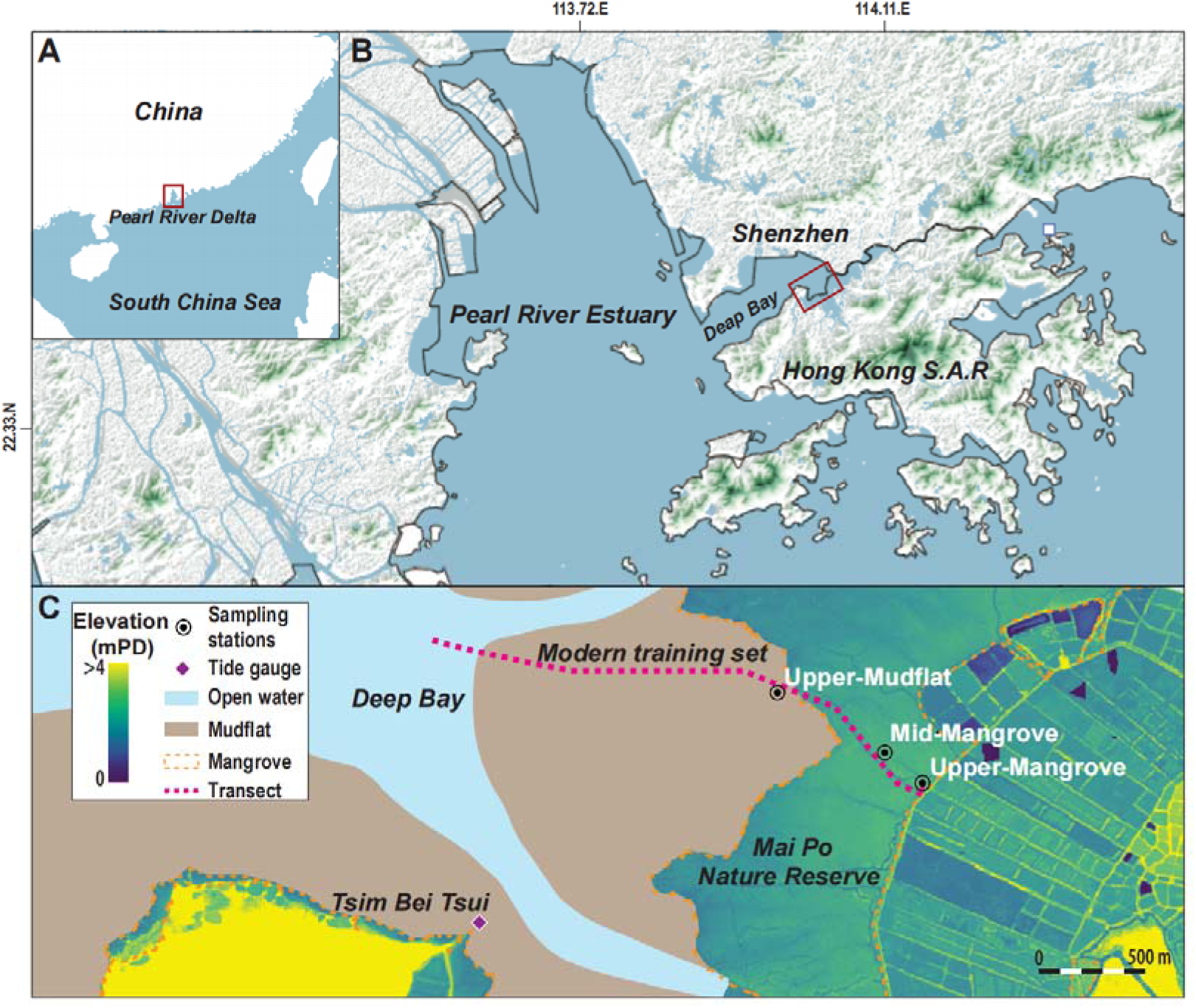
Study area and sampling locations. Location of the study area in the Pearl River Delta (A) and Deep Bay (B). (C) Monitoring stations set up at interior mangrove forest (upper-mangrove station), mangrove forest (mid-mangrove station) and mudflat-mangrove transitional zone (upper-mudflat station) are shown, along with the nearest tide gauge at Tsim Bei Tsui. The position of surface transect of established modern training set in Liu et al. (2025) is also shown. Land surface elevation was derived from the LiDAR digital elevation model supplied by the Hong Kong S.A.R. government for the Mai Po areas.

The climate is humid subtropical, characterized by distinct wet and dry seasons driven by seasonal monsoons (Ding & Chan, 2005; Yu et al., 2023). The southwest monsoon brings the wet season (April-September) that accounts for 77% of annual precipitation, with maximum monthly totals over 1000 mm. The northeast monsoon corresponds with the dry season (October-March), when lowest monthly total precipitation drop below 1 mm (HongKong-Observatory, 2024) (Fig. 2; Table S1). The precipitation shift between dry and wet seasons typically lags about one month behind monsoon onset, with main rains beginning in late April to early May and ending from late October to early November (HongKong-Observatory, 2024). Average monthly air temperature during the study period (from November 2022 to May 2024) ranged from 15-30 °C, and was lowest in February (15 °C) and highest in July (30 °C). The seasonal contrast between wet and dry conditions affects porewater pH, salinity, and other environmental factors (Chen et al., 2022; Li et al., 2011; Wang et al., 2013), likely influencing foraminiferal community composition and eDNA detection.

**Figure 2.**
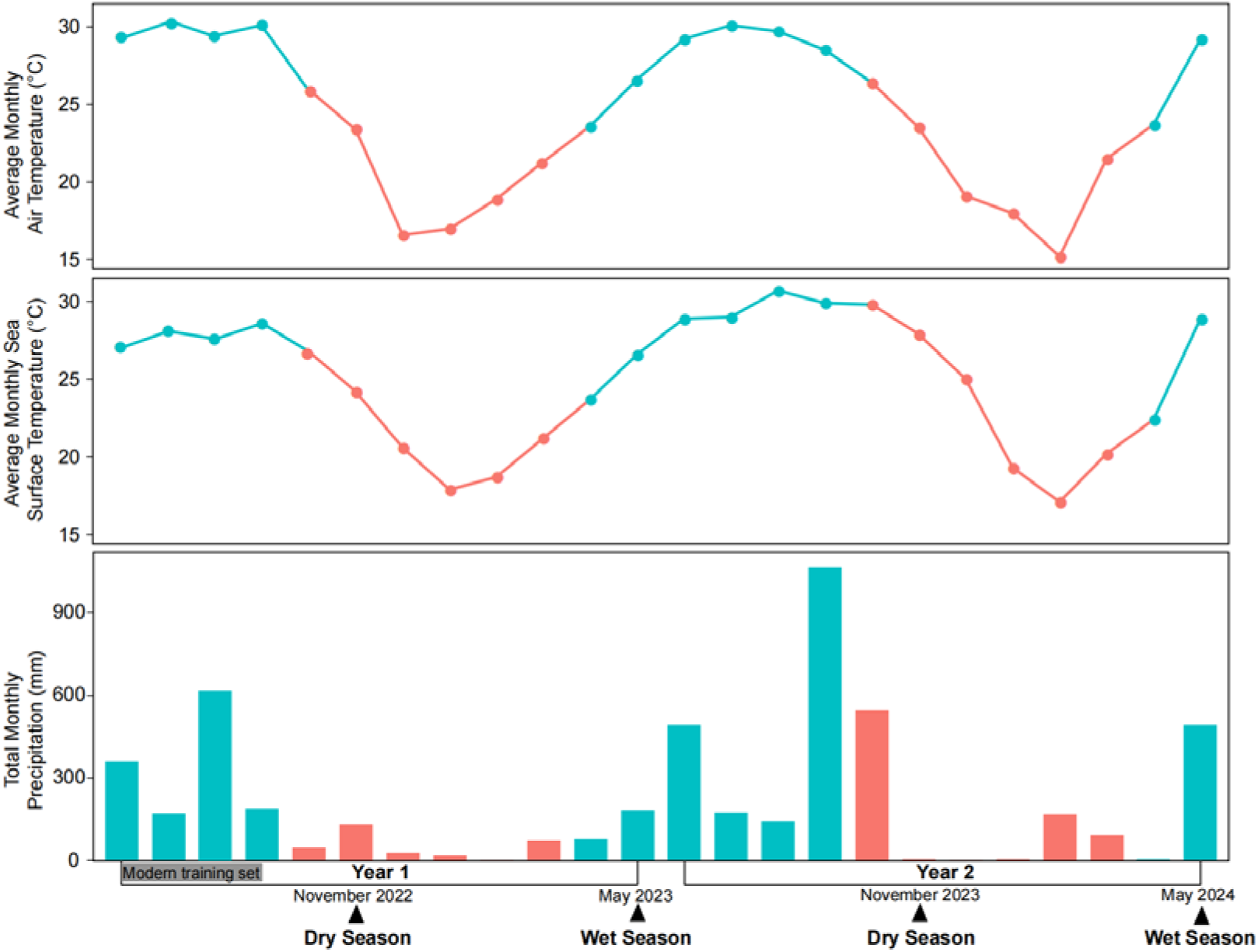
Climatic conditions at the study area during the monitoring period (Nov 2022-May 2024), with the sampling period for modern training set samples from Liu et al. (2025) also indicated. Time series of monthly averaged air temperature (top panel), sea surface temperature (middle panel), and total precipitation (bottom panel) show characteristic dry (red) and wet (blue) seasonal climate patterns of the region. Monthly averaged air temperature and total precipitation data were obtained from the monitoring station at Lau Fau Shan (the nearest station to our study area), while sea surface temperature data were sourced from the monitoring station at Hong Kong International Airport, both maintained by the Hong Kong Observatory (HongKong-Observatory, 2024).

Within Mai Po, a shore-normal transect was established spanning the mudflat-mangrove transitional zone to the interior mangrove forest, coinciding with a previous foraminiferal eDNA modern training set (Fig. 1C) (Liu et al., 2025). This transect captures the ecological gradient from baren mudflat to mature mangrove forest, enabling assessment of how environmental factors shape foraminiferal eDNA assemblage. Three monitoring stations were positioned along this transect: (1) mudflat-mangrove transitional zone (upper-mudflat station), (2) mangrove forest (mid-mangrove station), and (3) interior mangrove forest (upper-mangrove station) (Fig. 1C), to capture the temporal and small-scale variation of eDNA assemblages along the seaward-landward gradient. The flora in mudflat-mangrove transitional zone is dominated by *Sporobolus alterniflorus*, with occasional non-native *Sonneratia apetala* (Yu et al., 2025), while the mangrove stands are primarily occupied by *Kandelia candel*, *Aegiceras corniculatum*, and *Avicennia marina* (Lee, 2000; Li et al., 2019).

### 2.2 Sampling strategy

Given the study area’s distinct dry and wet season pattern, we conducted semi-annual sampling during the dry and wet seasons. Samples were collected at each monitoring station in late November (dry season) and late May (wet season), after the seasonal climatic factors (e.g., precipitation) had already shifted for at least two weeks (e.g. the first dry season collection occurred two weeks after the main precipitation had ceased by November 3rd, which accounted for >70% of that month’s total rainfall; The main wet-season rainfall in 2023 began on the 5^th^ May, two weeks before our collection period) (HongKong-Observatory, 2024). This timing ensured that the foraminiferal eDNA assemblages captured reflected true seasonal variation related to the respective collection periods. Within a 1 × 1 m^²^ plot at each monitoring station, three replicate sediment samples (approximately 20 cm^3^ each) for eDNA analysis were collected randomly from the top 1cm of the surface (Liu et al., 2025). This replication was designed to assess microscale spatial variability (patchiness) in foraminiferal eDNA assemblage composition within each monitoring station (Lejzerowicz et al., 2014; Walker et al., 2020). The position of each collected sample was recorded within 10 × 10 cm^2^ cells using a quadrat (Smith et al., 2021), to avoid repeatedly sampling from the same location across seasons. Sampling procedure followed Liu et al. (2025). Briefly, all sampling equipment was sanitized with 10% bleach between each collection to prevent cross-contamination. All eDNA samples were kept on ice during field collection and promptly transferred to a −20 °C freezer upon return to the laboratory on the sampling day.

To examine the potential environmental drivers of temporal variation in foraminiferal eDNA, an additional 20 cm^3^ of sediment was sampled simultaneously at each station to measure environmental variables, including porewater salinity and pH, and stable carbon isotope geochemistry [total organic carbon (TOC), total nitrogen (TN) and δ^13^C]. These samples were also kept on ice once collected and stored in a 4°C refrigerator in the lab.

The elevation of each monitoring station was measured following procedure introduced in Liu et al. (2025), relative to the Hong Kong Principal Datum (PD). Specifically, we took three elevation measurements at each monitoring station due to the slightly uneven surface topography. We converted the elevation to standardized water level index units (SWLI) (Kemp & Telford, 2015), where a value of 100 corresponds to mean tidal level (MTL) and 200 to mean higher high water (MHHW), based on tidal data obtained from the Tsim Bei Tsui tide gauge (Fig. 1C).

### 2.3 eDNA analysis

#### 2.3.1 eDNA extraction, amplification, and sequencing

We extracted eDNA from sediment samples using the 0.5-g PowerSoil® DNA Isolation Kit (QIAGEN). A two-step polymerase chain reaction (PCR) amplification protocol was conducted to prepare sequencing libraries. To minimize PCR bias and to cover total genetic/taxonomic diversity (Nichols et al., 2018; Singer et al., 2023), DNA extraction was performed in 2 replicates for each sample, followed by duplicate PCR amplifications of each extraction replicate. The PCR amplification procedure follows Liu et al. (2025). Briefly, the first PCR was conducted with primers s14F1-s15 to amplify the 135-190 bp fragment within the hypervariable region 37f of foraminiferal 18s SSU rDNA gene (Pawlowski et al., 2014). Both primers included overhang adapter sequences compatible with Illumina indices following Illumina MiSeq System sequencing protocol. Replicate PCR products from each sample were pooled and purified using AMPpure XP beads (Beckman Coulter, Singapore) following Illumina MiSeq System sequencing protocol. Subsequently, an indexing PCR was performed using the Nextera XT Index Kit (Illumina) to append dual indices to the purified amplicons. Indexed PCR products were purified again using AMPpure XP beads. Purified samples were pooled to a final concentration of 20μM in 10 mM Tris (pH 8.5) and sent to Novogene Co, Ltd for sequencing on the NovaSeq platform using paired-end 150 bp reads (PE 150).

#### 2.3.2 Bioinformatics

Bioinformatics processing followed the pipeline introduced in Liu et al. (2025). Briefly, raw sequence data were demultiplexed and primers removed by Cutadapt v2.8 (Martin, 2011), produced pair-end reads were further merged and trimmed by PEAR v0.9.11 with a quality threshold of 26 (Zhang et al., 2014). Concatenated sequences were denoised and chimeras removed using VSEARCH (Rognes et al., 2016). Sequences were clustered into operational taxonomic units (OTUs) at 97% similarity using VSEARCH. Quality control was applied to discard OTUs that were either shorter than 90 bp/longer than 230 bp, had fewer than 10 reads, or appeared in negative controls with read counts equal to or exceeding those in any sample. Taxonomy was assigned using BLASTn v2.12.0 (Altschul et al., 1990) with e-value of 0.001 at species, family, and phylum levels based on identity thresholds of 98%, 90%, and 80%, respectively, referring to GenBank nucleotide database and an in-house foraminiferal DNA database (Table S2). OTUs with identities between 80% and 90% (assigned to phylum level) and OTUs with unknown taxonomy were classified as “undetermined OTUs” and treated individually as taxa in subsequent analyses. OTUs with identity thresholds below 80% were not considered foraminifera and were removed.

### 2.4 Environmental variables

To assess the influence of environmental variables on eDNA assemblage composition over time, porewater salinity and pH, and sediment total organic carbon (TOC), total nitrogen (TN) and δ^13^C, were measured following the procedures introduced in Liu et al. (2025). Briefly, salinity and pH were measured using a Thermo Scientific™ A3255 pH/conductivity multimeter from porewater obtained by centrifuging sediment samples (Sawai et al., 2016). Sediment samples for δ^13^C, TOC, and C/N analysis were pretreated with 5% hydrochloric acid (HCl) prior to analysis on a EuroVector EA3028 Nu Horizon isotope-ratio mass spectrometer (IRMS) and EA Isolink Elemental Analyser. Additionally, climate data (air temperature, sea surface temperature and precipitation) during our study period (November 2022-May 2024) were obtained from the monitoring station at Lau Fau Shan (nearest to our study area), with additional sea surface temperature measurements obtained from the Hong Kong International Airport station maintained by the Hong Kong Observatory (HongKong-Observatory, 2024) to evaluate seasonal climatic influences on environmental variables and eDNA assemblages (Table S1).

### 2.5 Statistical analysis

To assess seasonal variation in the abundance of dominant foraminiferal taxa at each sampling station, we performed a one-way analysis of variance (ANOVA) with 1,000 permutations (p-value threshold = 0.05) following previous practices (Chung et al., 2024; Troth et al., 2021) (Table S3). Identical seasons from years 1 and 2 were merged as a single factor in this analysis. Dominant taxa were defined as those with >5% relative abundance in at least one sample. Prior to ANOVA, we assessed homogeneity of variances for each taxon’s abundance at each station and applied log-transformation as needed to meet ANOVA assumptions. The relative abundances of dominant taxa were plotted. To visualize patterns in taxonomic composition and diversity across environments and seasons, we generated a heatmap of dominant taxa by merging replicate samples at each station and season. This heatmap was produced using the ggheatmap function from the heatmaply package (Galili et al., 2018) in R (v4.4.2).

To examine overall patterns of temporal (seasonal) and spatial variation in foraminiferal eDNA assemblages, we conducted non-metric multidimensional scaling (NMDS) based on Bray-Curtis dissimilarity of taxa relative abundance data, using the vegan package (v2.6-4) in R (Oksanen et al., 2007). The NMDS ordination was constructed using dominant taxa (>5% relative abundance in at least one sample), and the first two NMDS dimensions were used to visualize compositional dissimilarities among stations and seasons. To test the significance of observed patterns, we used permutational multivariate analysis of variance (PERMANOVA) with 999 permutations (Anderson, 2014), also based on Bray-Curtis distances. PERMANOVA was used to evaluate whether eDNA assemblages differed significantly between seasons (with identical seasons from years 1 and 2 merged as a single factor) and among stations (p-value threshold = 0.05).

To evaluate microscale spatial variability (patchiness) within each monitoring station, we calculated pairwise Bray-Curtis dissimilarity among the three replicate samples collected within each 1 × 1 m² plot at every station and sampling event. The magnitude of within-station dissimilarity (differences between replicate samples collected simultaneously from the same monitoring station representing microscale spatial variability) was then compared to among-station dissimilarity (differences among samples collected during the same season from different monitoring stations) using a permutation test (10,000 permutations; p-value threshold = 0.05) (Welch, 1990). This analysis tested whether microscale patchiness within stations was significant relative to broader environmental differences. Additionally, ANOVA (1,000 permutations; p-value threshold = 0.05) was used to assess whether the dispersion (i.e., within-station variability) of eDNA assemblages differed significantly among the three monitoring stations during the same sampling period.

To determine the influence of environmental variables on the composition of foraminiferal eDNA assemblages, we performed redundancy analysis (RDA) and partial RDA (pRDA) using Hellinger-transformed relative abundance data, implemented in the vegan package (v2.6-4) in R (Legendre & Gallagher, 2001). Only dominant taxa (>5% relative abundance in at least one sample) were included in the RDA, consistent with the criteria for taxa applied in RSL reconstruction using a Bayesian transfer function (BTF) (Liu et al., 2025). The environmental variables considered included salinity, pH, TOC, carbon-nitrogen ratio (C/N), and δ^13^C. The significance (*p* <0.05) of environmental variables and the first two ordination axes were determined using ANOVA with 1,000 permutations. Additionally, linear mixed-effects models (LMMs) were performed to examine the significance of variation (p-value threshold = 0.05) in the environmental variables across seasons at each monitoring station (Luke, 2017). pRDA was conducted with tidal elevation (SWLI) of each monitoring station included. To further quantify the proportion of variance in assemblage composition explained by physicochemical (tidal elevation and environmental variables) and climatic (temperature, sea-surface temperature, and precipitation) variables, we conducted variance partitioning using the varpart function in vegan (Peres-Neto et al., 2006).

To investigate how seasonal variations in eDNA assemblages at each monitoring station potentially influenced sea-level reconstruction, we applied a BTF to estimate the elevation for each season at each station (Cahill et al., 2016). The BTF was constructed using the modern training set described in Liu et al. (2025), following the same processing procedure. Briefly, eDNA sequence counts were rarefied to the lowest sequencing depth in the dataset—after excluding one modern sample with low read numbers—to account for differences in sequencing depth among samples. The distribution of estimated elevations (SWLI) for each station and season was compared to observed elevations, and uncertainties were reported as the maximum value of the 2σ interval across replicates. Attempts to remove planktonic taxa Family Globigerinitidae to reduce uncertainty resulted in elevated overprediction at the upper-mudflat station and were therefore abandoned (Fig. S1).

## 3. Results

### 3.1 Seasonal Variation in Dominant Taxa

Several dominant foraminiferal taxa exhibited clear and, in some cases, statistically significant seasonal patterns in abundance across the three sampling stations (Fig. 3, 4; Table S2). For example, Saccamminidae were significantly more abundant during the dry season at the upper-mangrove and upper-mudflat stations (ANOVA; *p* >0.05), while Ammoniidae showed significantly higher abundance in the wet season at the mid-mangrove station. Detailed relative abundances for all taxa at each station are provided in Table S4.

**Figure 3.**
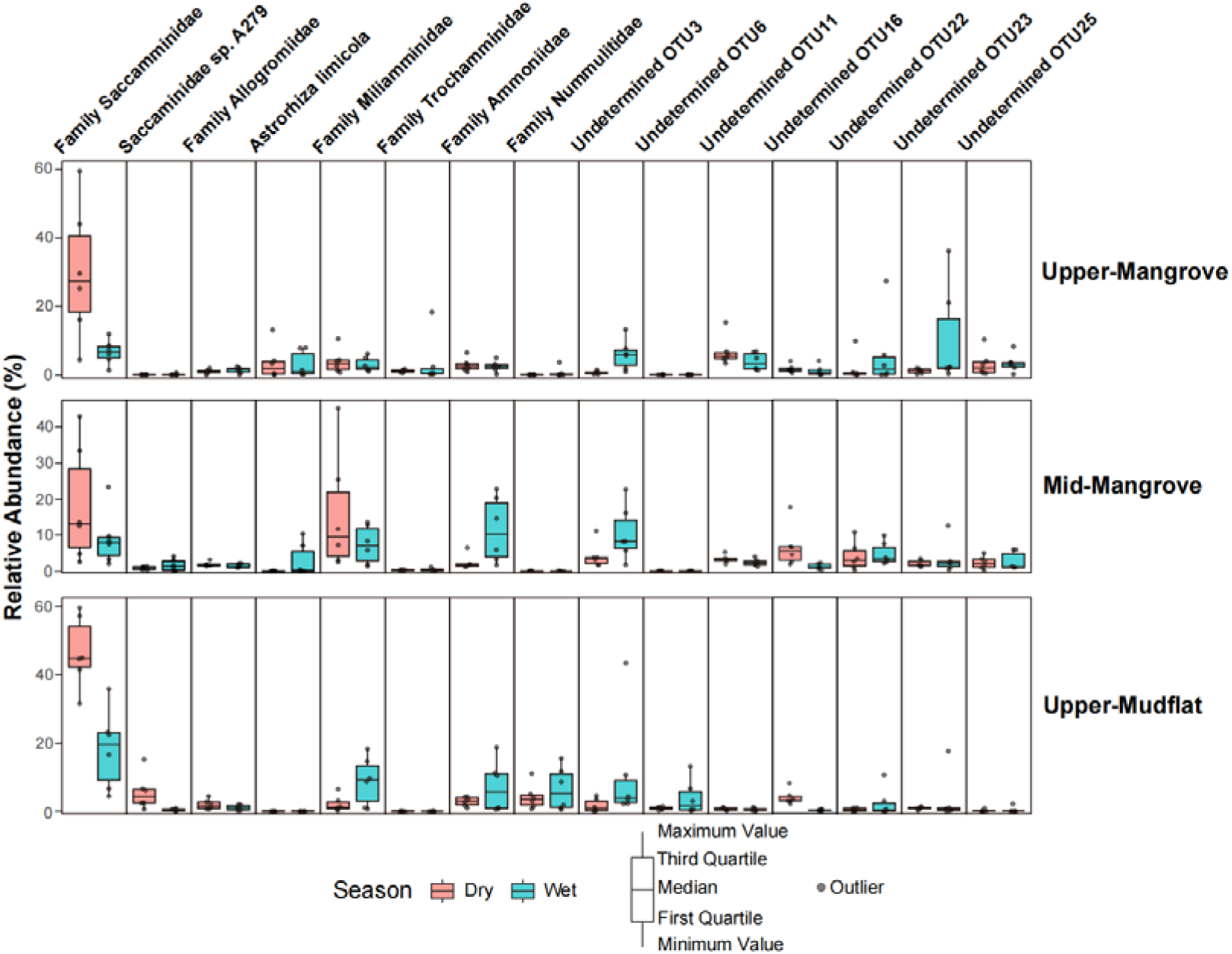
Relative abundance (%) of dominant foraminiferal taxa at three sampling stations during the wet (blue) and dry (red) seasons.

**Figure 4.**
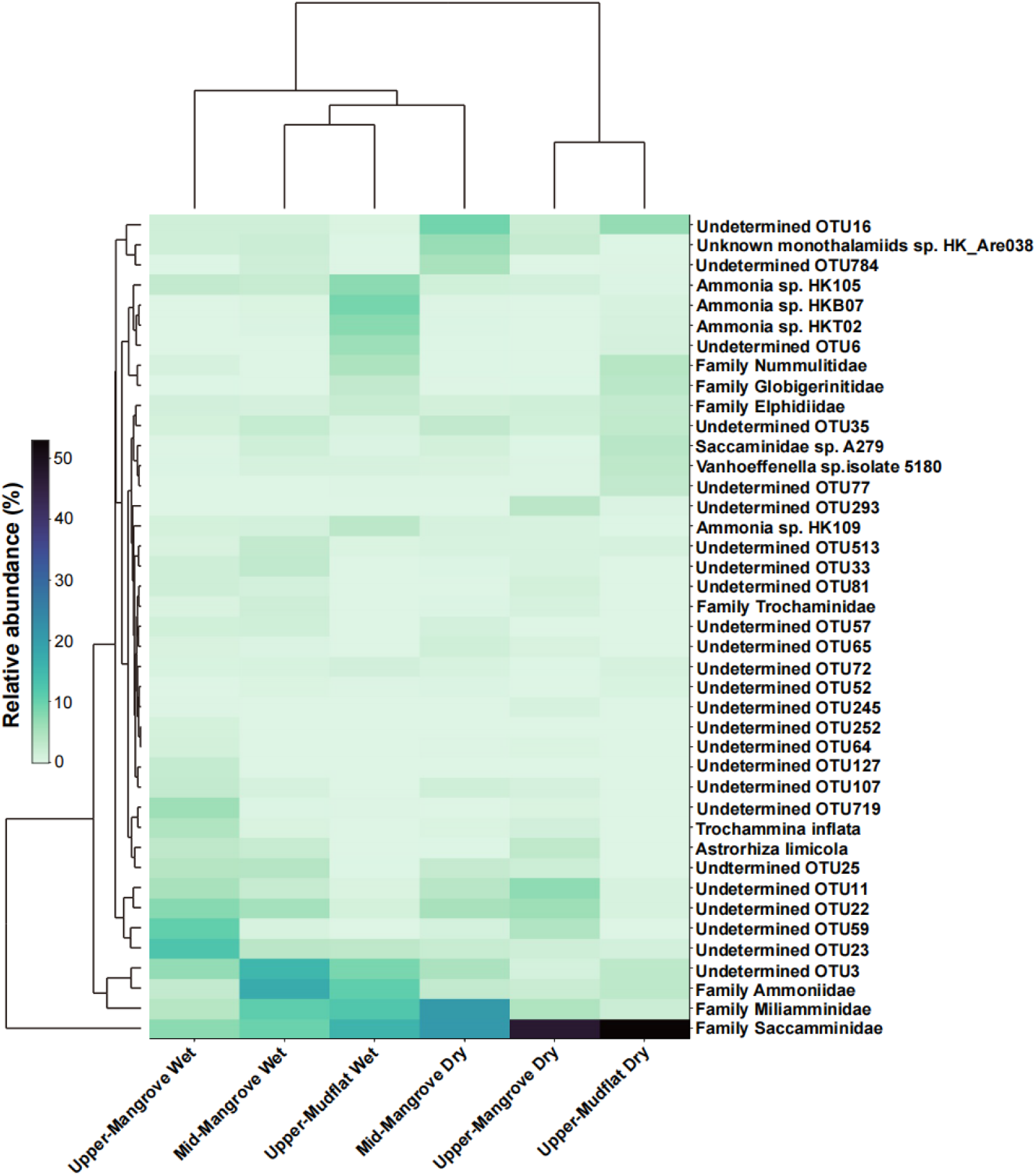
Heatmap of foraminiferal eDNA assemblages showing the relative abundance (%) of dominant foraminiferal taxa at each sampling station across wet and dry seasons.

### 3.2 Community structure across stations and seasons

The seasonal patterns in dominant taxa corresponded to distinct shifts in overall community composition across monitoring stations. During the dry season, the upper-mangrove, mid-mangrove, and upper-mudflat stations were all dominated by Saccamminidae, with Miliamminidae were also abundant in the mid-mangrove (Fig. 4). In the wet season, the upper-mangrove was dominated by OTU23 and OTU59, while the mid-mangrove was characterized by high abundances of Ammoniidae and OTU3. During the wet season, the upper-mudflat exhibited decreased abundance of Saccamminidae, but increased abundances of Miliamminidae, Ammoniidae, and OTU3 compared to the dry season (Fig. 4).

Environments as well as seasons influenced community compositions (Fig. 5A). Samples from the upper-and mid-mangrove clustered together, indicating that their foraminiferal community compositions were more similar to each other. In contrast, samples from the upper-mudflat formed a separate cluster, reflecting a distinct community composition (Fig. 5A). Within the upper-mangrove, dry and wet season samples largely overlapped in ordination space, yet PERMANOVA indicated a significant seasonal difference in community composition (F = 2.34, R² = 0.19, *p* = 0.01). In the upper-mudflat, dry and wet season samples were clearly separated in ordination space, also showing a significant difference between seasons (PERMANOVA; F = 4.42, R² = 0.31, *p* = 0.005). By contrast, no significant seasonal difference was detected in the mid-mangrove samples (PERMANOVA; F = 1.93, R² = 0.16, *p* = 0.079). In addition, most taxa were centrally located in the NMDS ordination, suggesting they were common to both the upper-mangrove and mid-mangrove environments. However, monothalamids such as Saccamminidae, and calcareous-walled Nummulitidae, were more closely associated with the upper-mudflat environment in the NMDS analysis (Fig. 5A).

**Figure 5.**
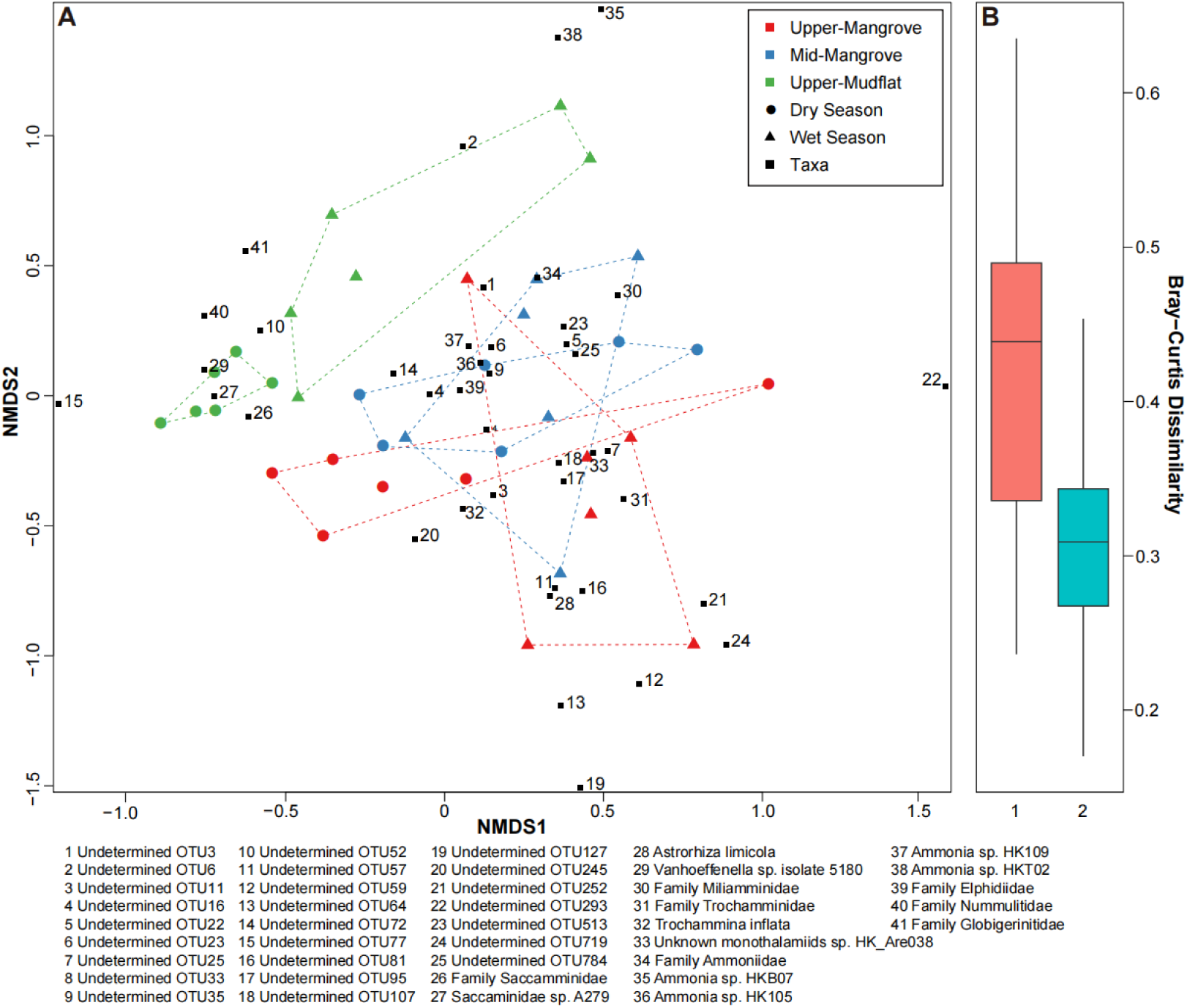
Dissimilarity of foraminiferal eDNA assemblage in each sample. (A) Non-metric Multi-Dimensional Scaling (NMDS) ordination of foraminiferal eDNA assemblages. NMDS ordination based on Bray-Curtis dissimilarities of foraminiferal eDNA assemblages at each sampling station during dry and wet seasons (stress value = 0.17). The NMDS was constructed using dominant taxa with >5% relative abundance in at least one sample. Samples from different environments and seasons are indicated by distinct colors and shapes. (B) Bar plot comparing average Bray-Curtis dissimilarity values among stations (1 = among-station variation) and within stations (2 = within-station variation).

### 3.3 Microscale (within-station) spatial variability

Samples showed significantly higher among-station heterogeneity than within-station (microscale) variability (Fig. 5B). Although the upper-mangrove station sampled in November 2022 exhibited the highest within-station dissimilarity (Table 1), within-station variability was significantly lower than among-station differences (permutation test; 10,000 permutations; *p* <0.01). Furthermore, overall variation in within-station dispersion across all monitoring stations was not statistically significant (ANOVA; *p* >0.05), indicating limited patchiness at the scale sampled.

**Table 1.** The average within-station Bray-Curtis (BC) Dissimilarity of each monitoring station per collection.

| Collection date | Monitoring station | Season | BC dissimilarity |
| --- | --- | --- | --- |
| 2022-November | Upper-mangrove | Dry | 0.46 |
|  | Mid-mangrove |  | 0.27 |
|  | Upper-mudflat |  | 0.26 |
| 2023-November | Upper-mangrove | Dry | 0.31 |
|  | Mid-mangrove |  | 0.32 |
|  | Upper-mudflat |  | 0.17 |
| 2023-May | Upper-mangrove | Wet | 0.45 |
|  | Mid-mangrove |  | 0.30 |
|  | Upper-mudflat |  | 0.34 |
| 2024-May | Upper-mangrove | Wet | 0.36 |
|  | Mid-mangrove |  | 0.24 |
|  | Upper-mudflat |  | 0.31 |

### 3.4 Influence of environmental variables

Both static (elevation) and seasonally variable environmental factors were measured to assess their influence on foraminiferal eDNA assemblages. The upper-(2.17-2.18 m PD; 189-190 SWLI) and mid-mangrove (2.17-2.22 m PD; 190-195 SWLI) monitoring stations shared similar elevations due to the flat topography of the mangrove forest, compared to the upper mudflat station (1.27-1.31 m PD; 93-99 SWLI). The upper-mangrove and mid-mangrove environments are characterized by lower salinity and higher pH, while the upper-mudflat exhibits lower total organic carbon (TOC) and higher δ^13^C values (Fig. 6C; Table 2).

**Figure 6.**
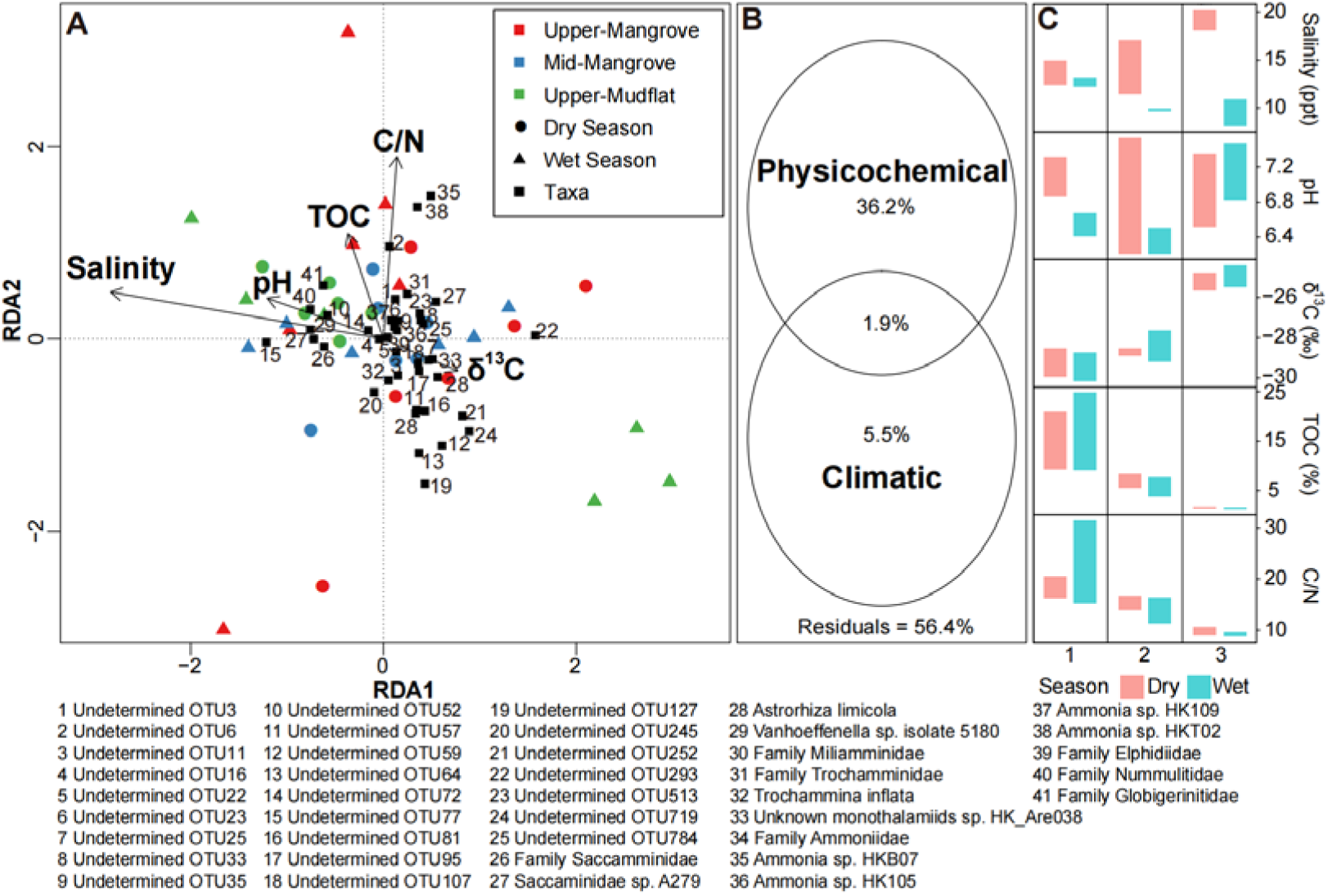
Environmental drivers of foraminiferal community structure. (A) Redundancy analysis (RDA) bioplot of dominant foraminiferal taxa (>5% relative abundance in at least one sample) and examined environmental variables. (B) Variance partitioning of community composition among physicochemical environmental variables and climatic data. Residual values are also shown. (C) Bar plots showing environmental variables measured at three monitoring stations during dry and wet seasons. (1 = upper-mangrove station, 2 = mid-mangrove station, 3 = upper-mudflat station).

**Table 2.**
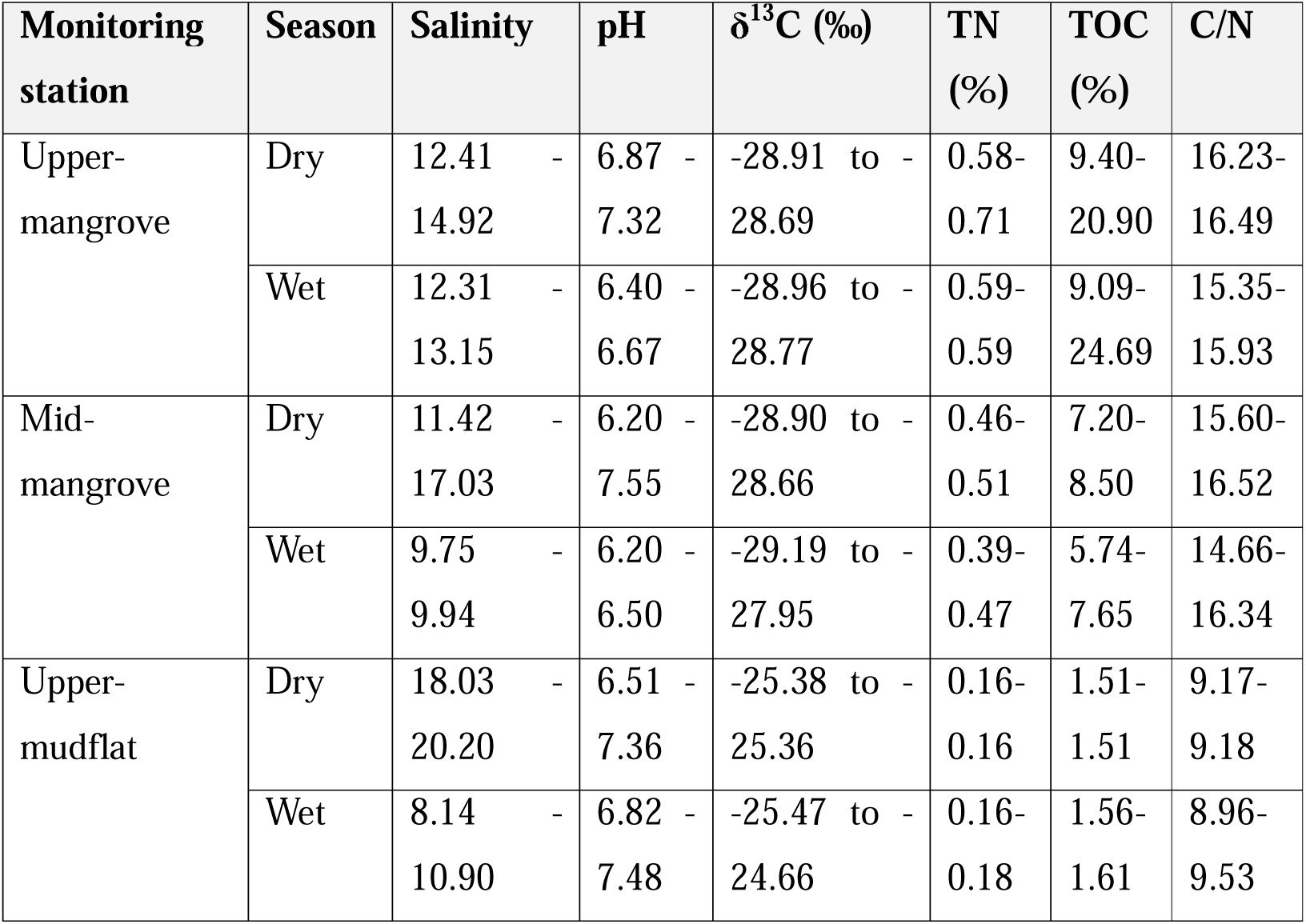
Environmental variables measured at each monitoring station per season.

Several measured environmental variables exhibited seasonal variation across the three environments (Fig. 6C; Table 2). Nevertheless, results from the LMMs indicated that only salinity at the upper-mudflat station exhibited a significant difference, being much higher in the dry season (18.0–20.2) than in the wet (8.1–10.9; LMMs; *p* <0.05). Taxa distribution was influenced by examined environmental conditions and climatic factors (Fig. 6). Salinity, pH, and TOC (*p* <0.01) significantly influenced foraminiferal eDNA composition across environments and seasons. Nevertheless, tidal elevation of each monitoring station remained the dominant driver, explaining 13% of the variance in eDNA assemblages, followed by salinity (11%) and pH (7%). Most taxa clustered near the center of the RDA bioplot, suggesting broad environmental tolerance, but some (e.g., Nummulitidae and Globigerinitidae) exhibited positive correlations with salinity and pH (Fig. 6A). Many undetermined OTUs (e.g., OTU59 and OTU127) showed strong negative correlations with TOC (Fig. 6A). Variance partitioning analysis revealed that physicochemical environmental properties explained 36.2% of the variance in eDNA assemblages, while climatic variables (including temperature, sea surface temperature and precipitation) accounted for 5.5% (Fig. 6B).

### 3.5 Relative sea level (RSL) estimation of each monitoring station

A foraminiferal eDNA BTF developed from previously established modern training set (Liu et al., 2025) was used with the eDNA dataset to estimate elevation for all replicate samples collected from each monitoring station across seasons and years. For most stations and years, the estimated elevations (within the 95% uncertainty interval) fell within the observed elevation range for that station, except for the upper-mudflat station during the wet season (Fig. 7). At this station, the BTF consistently overpredicted elevations in both seasons. The wet season samples from the upper-mangrove station exhibited the narrowest prediction range (187–198 SWLI), while the wet season samples from the upper-mudflat station had the lowest average 1-sigma uncertainty (9.6 SWLI; 0.48 m) (Table S5).

**Figure 7.**
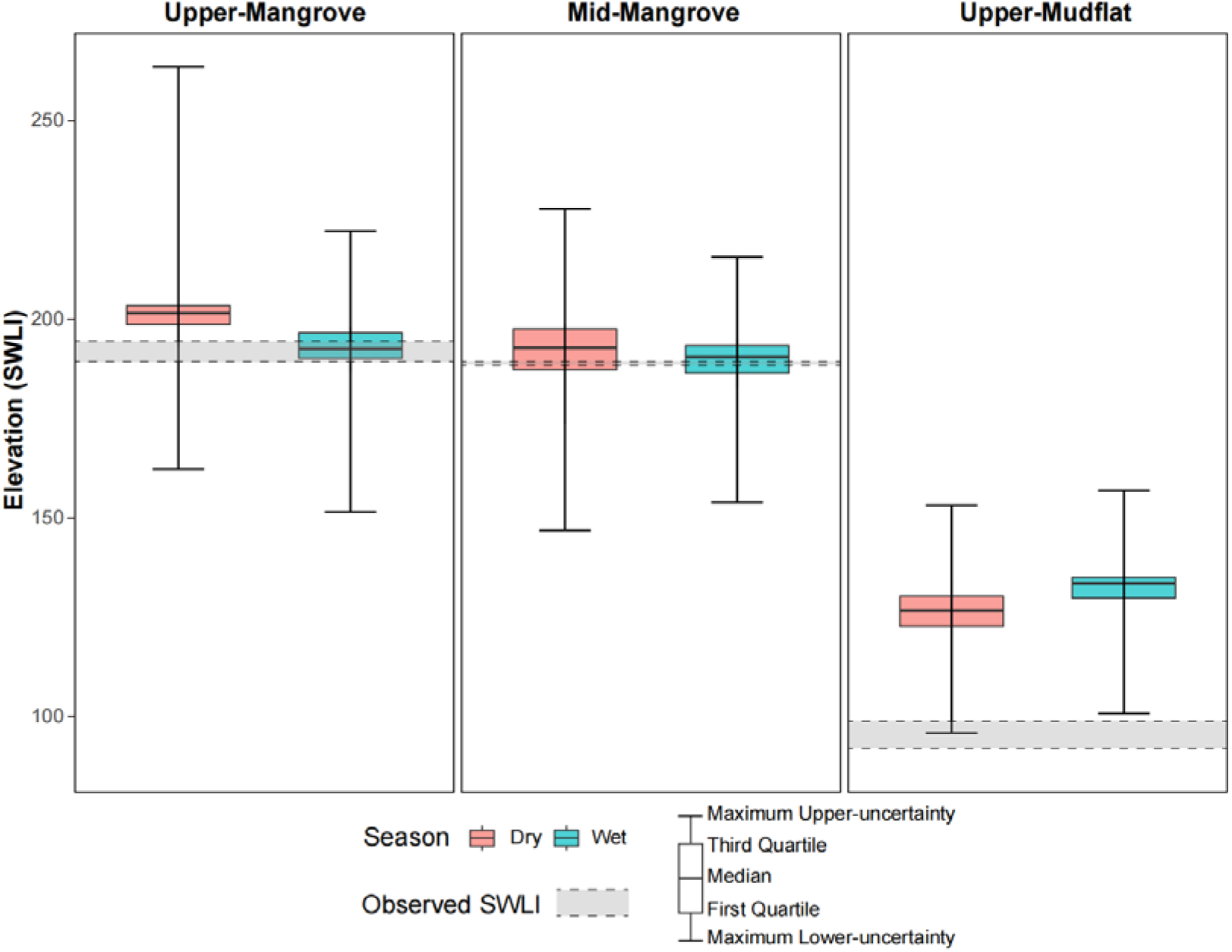
Elevation estimates based on foraminiferal eDNA assemblage. Elevation estimates (Standardized Water Level Index unit, SWLI) for each monitoring station during dry and wet seasons. Distributions represent six replicate samples per station and season. Error bars indicate the maximum 2σ uncertainty across replicates. The range of observed elevation at each station is shown by the grey dash bar.

## 4. Discussion

### 4.1 Temporal and spatial variation of foraminiferal eDNA assemblage

Foraminiferal eDNA assemblages varied temporally across seasons within specific habitats, but were spatially stable at small scales (i.e., low patchiness) across our three intertidal monitoring stations. Temporal variation patterns likely reflected fundamental differences in the composition and sources of eDNA across environments. At the mid-mangrove station, community composition remained relatively stable throughout the year. This stability suggests that this environment may buffer short-term seasonal environmental changes—as observed by previous studies in mangroves (Alongi, 2022; Camp et al., 2016)—or that its eDNA pool is dominated by accumulated extracellular eDNA from local populations across multiple generations, which can contribute >40% of total sediment eDNA (Ascher et al., 2009; Carini et al., 2016; Caro et al., 2023). This interpretation—that extracellular DNA may potentially dominate the assemblage—is consistent with traditional morphological studies of foraminifera in intertidal zones, where dead foraminiferal assemblages also show little seasonal variability compared to living (rose Bengal stained) assemblages (Buzas et al., 2015; Hippensteel et al., 2002; Horton & Edwards, 2003; Horton & Murray, 2006; Walker et al., 2020). Such stable, time-integrated foraminiferal communities—identified both in morphological analyses of dead assemblages and in eDNA assemblages dominated by autochthonous extracellular eDNA—tend to reflect long-term environmental gradients, such as tidal elevation, which are shaped by persistent factors like the duration and frequency of inundation or subaerial exposure (Horton & Culver, 2008; Woodroffe et al., 2005).

In contrast, the upper-mangrove and upper-mudflat stations exhibited marked seasonal changes in the relative abundance of dominant taxa such as Saccamminidae and OTU3 (Fig. 3, 4). This indicates a greater contribution from living populations in these environments, making the eDNA assemblages more responsive to short-term environmental fluctuations (Ellegaard et al., 2020; Goldberg et al., 2011; Singer et al., 2023; Torti et al., 2018). This pattern, commonly observed in eDNA studies conducted with modern samples (e.g. surface soils/marine water) (Angeles et al., 2020; Mazurkiewicz et al., 2024; Nagler et al., 2022), suggests that when DNA from living populations dominates over extracellular sources, assemblages capture more dynamic, short-term environmental changes—resulting in pronounced seasonal variability (Yun et al., 2023).

Ecologically, the observed seasonal variation in eDNA assemblages among stations appear to be driven by the life histories and environmental sensitivities of key foraminiferal taxa. Hard-shelled foraminifera, such as those in the family Ammoniidae and several of its OTUs, exhibited significant seasonal variation (Table S3), with higher abundance in the wet season at the mid-mangrove station (Fig. 3). This is consistent with morphological surveys, which report that calcareous taxa typically reach peak densities during spring and summer, coinciding with warmer temperatures and increased phytoplankton and zooplankton productivity (Alve & Murray, 1999; Berkeley et al., 2008; Horton & Edwards, 2003; Murray & Alve, 2000; Richirt et al., 2020). These seasonal population cycles likely lead to increased eDNA production rates during reproductive periods (Buxton et al., 2017; Fukumoto et al., 2015; Spear et al., 2015), resulting in changes of eDNA assemblages that respond more rapidly than morphological methods and influence the taxonomic composition detected in our samples. In contrast, monothalamous taxa—particularly Saccamminidae—were especially abundant during the dry season and contributed disproportionately to the observed seasonality. These taxa are known to be more sensitive to environmental change (He et al., 2019), and their seasonal dynamics are not well captured in traditional morphological surveys due to poor preservation of their delicate tests with the routine methodology (Schönfeld et al., 2012). Recent studies suggest that Saccamminidae may decline during summer, possibly due to competition with algal blooms (Henderson, 2023), while eDNA surveys in French estuarine mudflats have reported their highest abundances in the fall (Singer et al., 2023). This pronounced seasonality in Saccamminidae abundance helps explain our earlier RSL reconstruction efforts at this site (Liu et al., 2025), where the modern training set—based on wet season samples (primarily collected between June and August 2022)—showed comparatively lower Saccamminidae representation. In that study, Saccamminidae abundances reached up to 23% (from MTL to MHHW), similar to the 9–36% observed in the wet season samples of the present study (Fig. 3), but much lower than the abundances observed during the dry season (4-60%; Fig. 3). Although including monothalamids improve BTF performance (Liu et al., 2025), our results suggest that the timing of the collection of training set samples is an important consideration. In our study area, sample collection during the wet season—when Saccamminidae are less dominant—yields more balanced representation of indicator taxa and prevent compositional bias in eDNA data (Gloor et al., 2017). This approach is preferable for new studies aiming to explore the relationship between eDNA assemblages and tidal elevation, as it reduces the risk of dominant taxa masking underlying ecological patterns.

Our study also provides insights into the spatial variability of eDNA assemblages. In RSL reconstruction, small-scale spatial variability in foraminiferal assemblages may introduce bias to modern training sets, reducing their accuracy in capturing tidal elevation signals (Kemp et al., 2011). The upper-mangrove station exhibited the greatest heterogeneity among replicate samples (Table 1), but overall community structure and composition of dominant taxa remained consistent across all monitoring stations, indicating strong spatial stability at microscales (within 1 x 1 m^2^). Although the small-scale spatial variability in intertidal foraminiferal communities can be driven by the adaptation of living populations to local microhabitats and biotic factors such as predation pressure (Buzas, 1970, 1982) or food resource availability (Alve & Murray, 2001; Fontanier et al., 2003), the environments and conditions at our study site seemed to be relatively homogeneous. Consequently, the eDNA assemblages were primarily shaped by larger-scale environmental differences among different environments (Fig. 6C), and did not display the significant spatial patchiness observed in other environments, such as in the deep-sea (Lejzerowicz et al., 2014) or temperate mudflat environments (Singer et al., 2023). Furthermore, compared to morphological methods, metabarcoding incorporates DNA source from propagules (Hervé et al., 2022; Pont et al., 2018; Yang et al., 2022), particularly in mudflat environments (Liu et al., 2025; Singer et al., 2023), the mobility of which may give it greater potential for homogenization and thus representing environment-wide signals at small scales. This spatial stability has important implications for both paleoenvironmental reconstruction and contemporary monitoring. It supports the sampling design used in modern training sets for tidal elevation reconstruction, where single samples from narrow elevation intervals (typically <10 cm) are assumed representative of community structure at that elevation. It also suggests that eDNA-based monitoring in mangrove and mudflat systems can reliably capture the dominant community structure without intensive spatial replication under relatively uniform environmental conditions. This supports the use of single or pooled samples to represent local assemblages in coastal monitoring programs.

### 4.2 Factors influencing seasonality in foraminiferal eDNA assemblages

A range of abiotic and biotic environmental factors have long been recognized as important drivers of foraminiferal community composition in intertidal settings (Debenay et al., 2006; Du Châtelet et al., 2009a; Du Chatelet et al., 2009b; Lei et al., 2017; Murray, 1992). Our results further demonstrate that temporal fluctuations in environmental variables, shaped by seasonal and tidal cycles, are closely linked to the observed seasonality in foraminiferal eDNA assemblages. Salinity, pH, and total organic carbon (TOC) are significantly correlated with seasonal variation in eDNA community structure shown by RDA. These variables, in turn, are influenced by broader climatic drivers such as temperature, precipitation, and river discharge. Understanding these relationships is critical for the development and application of foraminiferal eDNA as an ecological indicator, as it highlights which environmental variables must be monitored or controlled to ensure reliable interpretation.

Among the short-term environmental variables examined (Table 2), salinity was the primary driver of seasonal changes in foraminiferal eDNA assemblages (Fig. 6A). Wet season precipitation and increased river discharge lowered salinity, favoring taxa adapted to reduced salinity (e.g., *Trochammina inflata*, which increased from 1–2% to 1–18%; Fig. 4), while constraining more stenohaline taxa intolerant of broader salinity ranges (Jorissen et al., 2022; Li et al., 2023; Singer et al., 2023). In parallel,, lower pH during the wet season likely influenced the abundance of calcareous foraminifera by affecting the calcification and metabolic processes (Dong et al., 2020; Le Cadre et al., 2003) (e.g. Nummulitidae; increased from <0.1% to up to 4%; Fig. 4, 6A), while fluctuations in TOC content indicated changes in food availability important for juvenile recruitment (Murray, 1992). Collectively, these variables shaped the distinct seasonal community patterns observed across the intertidal zone (Fig. 3, 4).

In addition to environmental factors like salinity and pH, climatic variables such as temperature also influence foraminiferal assemblages in intertidal zones, as shown by both morphological studies (Kaminski et al., 2021; Lei et al., 2017; Murray & Alve, 2000) and eDNA-based studies (Singer et al., 2023). However, variance partitioning in our study revealed that physicochemical environmental variables collectively explained 36.2% of variation in eDNA assemblages, while climatic factors collectively explained only 5.5% of the variation in eDNA assemblages (Fig. 6B). Although temperature can influence enzyme activity and, consequently, foraminiferal growth (Dong et al., 2019; Lombard et al., 2009; Prazeres & Pandolfi, 2016), the seasonal temperature variation in our study area per sampling was relatively small (up to 4.6 °C; Fig. 2; Table S1). Therefore, it is likely that climatic factors primarily exert indirect effects on foraminiferal communities by modulating local short-term environmental conditions.

Environmental variables not only affect living foraminiferal communities but may also influence the properties and preservation of eDNA itself. Higher temperatures during the wet season can accelerate extracellular eDNA degradation via increased physical stress and bacterial nuclease activity (Eichmiller et al., 2016; Lance et al., 2017; Strickler et al., 2015; Wood et al., 2020), potentially increasing the proportion of eDNA derived from living populations. Additionally, increased river discharge and stronger hydrodynamics during the wet season may elevate transportation of exogenous eDNA (e.g. propagules) into the intertidal zone (Harrison et al., 2019; Ji et al., 2020; Nevers et al., 2020), potentially introducing uncertainty when correlating tidal elevation with vertical zonation of foraminiferal eDNA assemblages.

Despite these seasonal effects, tidal elevation explained the largest proportion of variation in foraminiferal eDNA assemblages, highlighting its overriding and persistent influence that is consistent with previous morphological studies (Du Châtelet et al., 2018; Lei et al., 2017). Tidal elevation shapes distinct physicochemical gradients across the study area, as reflected in δ^13^C values and TOC content (Table 2), which both directly influence foraminiferal communities and mediate the effects of seasonal environmental changes at different elevations. While seasonal variations in salinity, pH, and TOC shape temporal patterns in foraminiferal assemblages, tidal elevation remains the dominant factor, emphasizing the key role of local environmental context in structuring intertidal communities.

### 4.3 Implications of temporal variation in foraminiferal eDNA for sea-level reconstruction

Our results demonstrate that foraminiferal eDNA assemblages serve as reliable proxies for tidal elevation in mangrove environments, with elevation estimates derived from eDNA BTFs for both dry and wet seasons typically falling within observed elevation ranges (Fig. 7). This reliability supports the use of mangrove-derived eDNA samples for constructing modern training sets throughout the year. In contrast to mangrove environments, the accuracy of eDNA-based reconstructions was reduced at the upper-mudflat station situated at a transitional zone, where elevation estimates were consistently overpredicted (Fig. 7). This limitation is likely driven by increased contributions of allochthonous and propagule-derived eDNA in these dynamic environments (Angeles et al., 2020; Goldstein & Alve, 2011; Singer et al., 2023), especially during the wet season. The potential influx of allochthonous eDNA, particularly from higher-elevation mangrove environments, expands the apparent environmental tolerances of some taxa (e.g., *Vanhoeffenella sp. isolate 5180*) and increases uncertainty in RSL reconstructions of transitional zone. Additionally, anthropogenic disturbance in transitional zones (Yu et al., 2025) may exacerbate habitat instability and further confound the eDNA signal. These findings mirror morphological studies, which report greater community turnover and weaker tidal elevation gradients at lower-elevation mudflats (Lei et al., 2017), resulting in larger offsets in RSL estimates.

Our findings support several recommendations for optimizing foraminiferal eDNA as a proxy for reconstructing RSL change: (1) prioritizing stable mangrove environments for constructing modern training sets, as these yield signals most closely linked to local tidal elevation; (2) avoiding environmental transitional zones, such as upper-mudflat where possible, or if necessary, sampling during the dry season following established morphological protocols (Horton & Edwards, 2003), to minimize allochthonous influences; and (3) utilizing single samples per homogeneous environment following morphological methods (Horton & Edwards, 2006; Kemp et al., 2009a; Kemp et al., 2012), which our results show are sufficient for spatial representation. By refining both sampling strategies and analytical approaches, foraminiferal eDNA has significant potential to provide more accurate and robust reconstructions of past RSL change in coastal wetlands.

## 5. Conclusion

Our study provides new insights into the temporal and spatial variation of foraminiferal eDNA assemblages and their implications for relative sea-level (RSL) reconstruction. Foraminiferal eDNA assemblage at the mid-mangrove monitoring stations remained relatively stable across seasons, demonstrating extracellular eDNA dominant, whereas the upper-mangrove station showed significant seasonal variation. The mudflat environment displayed distinctly different foraminiferal eDNA assemblages between the dry and wet seasons, showing higher exogenous eDNA contribution. Despite the influence of seasonally variable environmental and climatic factors, tidal elevation remained the dominant factor shaping eDNA assemblages. Elevation estimates produced by the eDNA BTF for the mangrove monitoring stations are accurate for both dry and wet seasons, while predictions for the upper-mudflat station during the wet season failed to be accurate. Our results reveal that foraminiferal eDNA assemblages display environment-specific seasonal variability, necessitating cautious interpretation of mudflat samples given their pronounced sensitivity to seasonal changes. Nevertheless, we demonstrate that eDNA remains a robust and broadly applicable indicator for RSL reconstruction, ensuring its utility for future studies.

## CRediT authorship contribution statement

**Liu Zhaojia**: Conceptualization, Data curation, Formal analysis, Methodology, Software, Visualization, Writing – original draft. **Nicole S. Khan**: Conceptualization, Funding acquisition, Project administration, Resources, Supervision, Writing – review and editing. **Howard K.Y. Yu**: Investigation, Writing – review and editing. **Magali Schweizer**: Methodology, Writing – review and editing. **Jennifer S. Walker**: Methodology, Writing – review and editing. **Celia Schunter**: Conceptualization, Resources, Supervision, Writing – review and editing.

## Supporting information

Supplementary Figure-Temporal and spatial variability of mudflat and mangrove foraminiferal eDNA assemblages and its implication for sea-level recon

Supplementary Table-Temporal and spatial variability of mudflat and mangrove foraminiferal eDNA assemblages and its implication for sea-level recons

## Acknowledgement

This project is funded by the Research Grants Council of Hong Kong (GRF Project no.: 27300221 and 17303925). We gratefully acknowledge the World Wide Fund for Nature for permitting this research in the Mai Po Nature Reserve and for their continuous conservation efforts in Hong Kong. We also extend our sincere appreciation to Mr. Gao Chengcheng and Miss Qin Yonghui (listed alphabetically) for their invaluable assistance during challenging fieldwork conditions.

## Appendix A. Supplementary data

Supplementary data of this article can be found at excel file *“Supplementary Table-Temporal and spatial variability of mudflat and mangrove foraminiferal eDNA assemblages and its implication for sea-level reconstruction”*.

## Data availability

Raw sequence data has been uploaded to the Sequence Read Archive under Bioproject: <u>PRJNA1306768</u>.

## References

1. Alongi, D. M. (2022). Climate change and mangroves. In Mangroves: biodiversity, livelihoods and conservation (pp. 175-198). Springer.

2. Altschul, S. F., Gish, W., Miller, W., Myers, E. W., & Lipman, D. J. (1990). Basic local alignment search tool. Journal of molecular biology, 215(3), 403–410. 10.1016/S0022-2836(05)80360-2

3. Alve, E., & Murray, J. W. (1999). Marginal marine environments of the Skagerrak and Kattegat: a baseline study of living (stained) benthic foraminiferal ecology. Palaeogeography, Palaeoclimatology, Palaeoecology, 146(1-4), 171–193. 10.1016/S0031-0182(98)00131-X

4. Alve, E., & Murray, J. W. (2001). Temporal variability in vertical distributions of live (stained) intertidal foraminifera, southern England. The Journal of Foraminiferal Research, 31(1), 12–24. 10.2113/0310012

5. Anderson, M. J. (2014). Permutational multivariate analysis of variance (PERMANOVA). Wiley statsref: statistics reference online, 1–15. 10.1002/9781118445112.stat07841

6. Angeles, I. B., Lejzerowicz, F., Cordier, T., Scheplitz, J., Kucera, M., Ariztegui, D., Pawlowski, J., & Morard, R. (2020). Planktonic foraminifera eDNA signature deposited on the seafloor remains preserved after burial in marine sediments. Scientific Reports, 10(1), 20351. 10.1038/s41598-020-77179-8

7. Ascher, J., Ceccherini, M. T., Pantani, O.-L., Agnelli, A., Borgogni, F., Guerri, G., Nannipieri, P., & Pietramellara, G. (2009). Sequential extraction and genetic fingerprinting of a forest soil metagenome. Applied Soil Ecology, 42(2), 176–181. 10.1016/j.apsoil.2009.03.005

8. Bakker, J., Wangensteen, O. S., Baillie, C., Buddo, D., Chapman, D. D., Gallagher, A. J., Guttridge, T. L., Hertler, H., & Mariani, S. (2019). Biodiversity assessment of tropical shelf eukaryotic communities via pelagic eDNA metabarcoding. Ecology and Evolution, 9(24), 14341–14355. 10.1002/ece3.5871

9. Barnett, R. L., Garneau, M., & Bernatchez, P. (2016). Salt-marsh sea-level indicators and transfer function development for the Magdalen Islands in the Gulf of St. Lawrence, Canada. Marine Micropaleontology, 122, 13–26. 10.1016/j.marmicro.2015.11.003

10. Berkeley, A., Perry, C., Smithers, S., & Horton, B. (2008). The spatial and vertical distribution of living (stained) benthic foraminifera from a tropical, intertidal environment, north Queensland, Australia. Marine Micropaleontology, 69(2), 240–261. 10.1016/j.marmicro.2008.08.002

11. Berkeley, A., Perry, C. T., & Smithers, S. G. (2009a). Taphonomic signatures and patterns of test degradation on tropical, intertidal benthic foraminifera. Marine Micropaleontology, 73(3-4), 148–163. 10.1016/j.marmicro.2009.08.002

12. Bista, I., Carvalho, G. R., Walsh, K., Seymour, M., Hajibabaei, M., Lallias, D., Christmas, M., & Creer, S. (2017). Annual time-series analysis of aqueous eDNA reveals ecologically relevant dynamics of lake ecosystem biodiversity. Nature communications, 8(1), 14087. 10.1038/ncomms14087

13. Bouchet, V. M., Frontalini, F., Francescangeli, F., Sauriau, P.-G., Geslin, E., Martins, M. V. A., Almogi-Labin, A., Avnaim-Katav, S., Di Bella, L., & Cearreta, A. (2021). Indicative value of benthic foraminifera for biomonitoring: Assignment to ecological groups of sensitivity to total organic carbon of species from European intertidal areas and transitional waters. Marine Pollution Bulletin, 164, 112071. 10.1016/j.marpolbul.2021.112071

14. Brinkmann, I., Schweizer, M., Singer, D., Quinchard, S., Barras, C., Bernhard, J. M., & Filipsson, H. L. (2023). Through the eDNA looking glass: Responses of fjord benthic foraminiferal communities to contrasting environmental conditions. Journal of Eukaryotic Microbiology, e12975. 10.1111/jeu.12975

15. Buragohain, D., & Ghosh, A. (2021). Seasonal Distribution Trends of Benthic Foraminiferal Assemblages from the Saurashtra Coast, Western India. Journal of the Geological Society of India, 97(1), 61–69. 10.1007/s12594-021-1626-1

16. Buxton, A. S., Groombridge, J. J., Zakaria, N. B., & Griffiths, R. A. (2017). Seasonal variation in environmental DNA in relation to population size and environmental factors. Scientific Reports, 7(1), 46294. 10.1038/srep46294

17. Buzas, M. A. (1970). Spatial homogeneity: statistical analyses of unispecies and multispecies populations of foraminifera. Ecology, 51(5), 874–879. 10.2307/1933980

18. Buzas, M. A. (1982). Regulation of foraminiferal densities by predation in the Indian River, Florida. Journal of Foraminiferal Research, 12(1), 66–71. 10.2113/gsjfr.12.1.66

19. Buzas, M. A., C Hayek, L.-A., Jett, J. A., & Reed, S. A. (2015). Pulsating patches: History and Analyses of spatial, seasonal, and yearly distribution of living benthic foraminifera (Vol. 97). Smithsonian Institution Scholarly Press.

20. Cahill, N., Kemp, A. C., Horton, B. P., & Parnell, A. C. (2016). A Bayesian hierarchical model for reconstructing relative sea level: from raw data to rates of change. Climate of the Past, 12(2), 525–542. 10.5194/cp-12-525-2016

21. Camp, E. F., Suggett, D. J., Gendron, G., Jompa, J., Manfrino, C., & Smith, D. J. (2016). Mangrove and seagrass beds provide different biogeochemical services for corals threatened by climate change. Frontiers in Marine Science, 3, 52. 10.3389/fmars.2016.00052

22. Carini, P., Marsden, P. J., Leff, J. W., Morgan, E. E., Strickland, M. S., & Fierer, N. (2016). Relic DNA is abundant in soil and obscures estimates of soil microbial diversity. Nature microbiology, 2(3), 1–6. 10.1038/nmicrobiol.2016.242

23. Caro, D. M., Horstmann, L., Ganzert, L., Oses, R., Friedl, T., & Wagner, D. (2023). An improved method for intracellular DNA (iDNA) recovery from terrestrial environments. MicrobiologyOpen, 12(3), e1369. 10.1002/mbo3.1369

24. Chen, J., Zhou, Z., & Gu, J.-D. (2022). Seasonal variations of n-damo bacterial community in the subtropical Mai Po mangrove wetland of Hong Kong. International Biodeterioration & Biodegradation, 175, 105503. 10.1016/j.ibiod.2022.105503

25. Chung, A., Kam, Y., Shea, S. K., & Schunter, C. (2024). Detecting Fish Diversity in Urban - Impacted Ecosystems: A Comparative Approach of eDNA Metabarcoding and UVC. Environmental DNA, 6(6), e70048. 10.1002/edn3.70048

26. Debenay, J.-P., Bicchi, E., Goubert, E., & Du Châtelet, E. A. (2006). Spatio-temporal distribution of benthic foraminifera in relation to estuarine dynamics (Vie estuary, Vendée, W France). Estuarine, Coastal and Shelf Science, 67(1-2), 181–197. 10.1016/j.ecss.2005.11.014

27. Ding, Y., & Chan, J. C. (2005). The East Asian summer monsoon: an overview. Meteorology and Atmospheric Physics, 89(1), 117–142. 10.1007/s00703-005-0125-z

28. Dong, S., Lei, Y., Li, T., & Jian, Z. (2019). Responses of benthic foraminifera to changes of temperature and salinity: Results from a laboratory culture experiment. Science China Earth Sciences, 62, 459–472. 10.1007/s11430-017-9269-3

29. Dong, S., Lei, Y., Li, T., & Jian, Z. (2020). Response of benthic foraminifera to pH changes: Community structure and morphological transformation studies from a microcosm experiment. Marine Micropaleontology, 156, 101819. 10.1016/j.marmicro.2019.101819

30. Du Châtelet, É. A., Bout-Roumazeilles, V., Riboulleau, A., & Trentesaux, A. (2009a). Sediment (grain size and clay mineralogy) and organic matter quality control on living benthic foraminifera. Revue de micropaléontologie, 52(1), 75–84. 10.1016/j.revmic.2008.10.002

31. Du Chatelet, E. A., Degre, D., Sauriau, P.-G., & Debenay, J.-P. (2009b). Distribution of living benthic foraminifera in relation with environmental variables within the Aiguillon cove (Atlantic coast, France): improving knowledge for paleoecological interpretation. Bulletin de la Société Géologique de France, 180(2), 131–144. 10.2113/gssgfbull.180.2.131

32. Du Châtelet, E. A., Francescangeli, F., Bouchet, V., & Frontalini, F. (2018). Benthic foraminifera in transitional environments in the English Channel and the southern North Sea: A proxy for regional-scale environmental and paleo-environmental characterisations. Marine Environmental Research, 137, 37–48. 10.1016/j.marenvres.2018.02.021

33. Edwards, R., & Horton, B. (2000). Reconstructing relative sea-level change using UK salt-marsh foraminifera. Marine Geology, 169(1-2), 41–56. 10.1016/S0025-3227(00)00078-5

34. Eichmiller, J. J., Best, S. E., & Sorensen, P. W. (2016). Effects of temperature and trophic state on degradation of environmental DNA in lake water. Environmental Science & Technology, 50(4), 1859–1867. 10.1021/acs.est.5b05672

35. Ellegaard, M., Clokie, M. R., Czypionka, T., Frisch, D., Godhe, A., Kremp, A., Letarov, A., McGenity, T. J., Ribeiro, S., & John Anderson, N. (2020). Dead or alive: sediment DNA archives as tools for tracking aquatic evolution and adaptation. Communications Biology, 3(1), 169. 10.1038/s42003-020-0899-z

36. Fontanier, C., Jorissen, F., Chaillou, G., David, C., Anschutz, P., & Lafon, V. (2003). Seasonal and interannual variability of benthic foraminiferal faunas at 550 m depth in the Bay of Biscay. Deep Sea Research Part I: Oceanographic Research Papers, 50(4), 457–494. 10.1016/S0967-0637(02)00167-X

37. Fouet, M. P., Schweizer, M., Singer, D., Richirt, J., Quinchard, S., & Jorissen, F. J. (2024). Unravelling the distribution of three Ammonia species (Foraminifera, Rhizaria) in French Atlantic Coast estuaries using morphological and metabarcoding approaches. Marine Micropaleontology, 188, 102353. 10.1016/j.marmicro.2024.102353

38. Frontalini, F., Buosi, C., Da Pelo, S., Coccioni, R., Cherchi, A., & Bucci, C. (2009). Benthic foraminifera as bio-indicators of trace element pollution in the heavily contaminated Santa Gilla lagoon (Cagliari, Italy). Marine Pollution Bulletin, 58(6), 858–877. 10.1016/j.marpolbul.2009.01.015

39. Fukumoto, S., Ushimaru, A., & Minamoto, T. (2015). A basin-scale application of environmental DNA assessment for rare endemic species and closely related exotic species in rivers: A case study of giant salamanders in Japan. Journal of applied ecology, 52(2), 358–365. 10.1111/1365-2664.12392

40. Galili, T., O’Callaghan, A., Sidi, J., & Sievert, C. (2018). heatmaply: an R package for creating interactive cluster heatmaps for online publishing. Bioinformatics, 34(9), 1600–1602. 10.1093/bioinformatics/btx657

41. Gloor, G. B., Macklaim, J. M., Pawlowsky-Glahn, V., & Egozcue, J. J. (2017). Microbiome datasets are compositional: and this is not optional. Frontiers in Microbiology, 8, 2224. 10.3389/fmicb.2017.02224/full

42. Goldberg, C. S., Pilliod, D. S., Arkle, R. S., & Waits, L. P. (2011). Molecular detection of vertebrates in stream water: a demonstration using Rocky Mountain tailed frogs and Idaho giant salamanders. Plos One, 6(7), e22746. 10.1371/journal.pone.0022746

43. Goldstein, S. T., & Alve, E. (2011). Experimental assembly of foraminiferal communities from coastal propagule banks. Marine Ecology Progress Series, 437, 1–11. 10.3354/meps09296

44. Gu, S., Deng, Y., Wang, P., Li, C., Shi, D., & Wang, S. (2023). Assessing riverine fish community diversity and stability by eDNA metabarcoding. Ecological Indicators, 157, 111222. 10.1016/j.ecolind.2023.111222

45. Harrison, J. B., Sunday, J. M., & Rogers, S. M. (2019). Predicting the fate of eDNA in the environment and implications for studying biodiversity. Proceedings of the Royal Society B, 286(1915), 20191409. 10.1098/rspb.2019.1409

46. Hawkes, A. D., Horton, B., Nelson, A., & Hill, D. (2010). The application of intertidal foraminifera to reconstruct coastal subsidence during the giant Cascadia earthquake of AD 1700 in Oregon, USA. Quaternary International, 221(1-2), 116–140. 10.1016/j.quaint.2009.09.019

47. Hayward, B. W., Grenfell, H., Cairns, G., & Smith, A. (1996). Environmental controls on benthic foraminiferal and thecamoebian associations in a New Zealand tidal inlet. The Journal of Foraminiferal Research, 26(2), 150–171. 10.2113/gsjfr.26.2.150

48. He, X., Sutherland, T. F., Pawlowski, J., & Abbott, C. L. (2019). Responses of foraminifera communities to aquaculture-derived organic enrichment as revealed by environmental DNA metabarcoding. Molecular Ecology, 28(5), 1138–1153. 10.1111/mec.15007

49. Henderson, Z. (2023). Soft-walled monothalamid foraminifera from the intertidal zones of the Lorn area, north-west Scotland. Journal of the Marine Biological Association of the United Kingdom, 103, e18. 10.1017/S002531542300006

50. Hervé, A., Domaizon, I., Baudoin, J.-M., Dejean, T., Gibert, P., Jean, P., Peroux, T., Raymond, J.-C., Valentini, A., & Vautier, M. (2022). Spatio-temporal variability of eDNA signal and its implication for fish monitoring in lakes. Plos One, 17(8), e0272660. 10.1371/journal.pone.0272660

51. Hippensteel, S. P., Martin, R. E., Nikitina, D., & Pizzuto, J. E. (2002). Interannual variation of marsh foraminiferal assemblages (Bombay Hook National Wildlife Refuge, Smyrna, DE): Do foraminiferal assemblages have a memory? The Journal of Foraminiferal Research, 32(2), 97–109. 10.2113/0320097

52. Hong Kong-Observatory. (2024). SUMMARY OF METEOROLOGICAL AND TIDAL OBSERVATIONS IN HONG KONG. https://www.hko.gov.hk/en/publica/pubsmo.htm

53. Horton, B. (1999). The distribution of contemporary intertidal foraminifera at Cowpen Marsh, Tees Estuary, UK: implications for studies of Holocene sea-level changes. Palaeogeography, Palaeoclimatology, Palaeoecology, 149(1-4), 127–149. 10.1016/S0031-0182(98)00197-7

54. Horton, B. P., & Culver, S. J. (2008). Modern intertidal foraminifera of the Outer Banks, North Carolina, USA, and their applicability for sea-level studies. Journal of coastal research, 24(5), 1110–1125. 10.2112/08A-0004.1

55. Horton, B. P., & Edwards, R. J. (2003). Seasonal distributions of foraminifera and their implications for sea-level studies, Cowpen Marsh, UK. 10.2110/pec.03.75.0021

56. Horton, B. P., & Edwards, R. J. (2006). Quantifying Holocene sea level change using intertidal foraminifera: lessons from the British Isles. Cushman Foundation for foraminiferal research.

57. Horton, B. P., & Murray, J. W. (2006). Patterns in cumulative increase in live and dead species from foraminiferal time series of Cowpen Marsh, Tees Estuary, UK: Implications for sea-level studies. Marine Micropaleontology, 58(4), 287–315. 10.1016/j.marmicro.2005.10.006

58. Ji, H., Pan, S., & Chen, S. (2020). Impact of river discharge on hydrodynamics and sedimentary processes at Yellow River Delta. Marine Geology, 425, 106210. 10.1016/j.margeo.2020.106210

59. Jorissen, F. J., Fouet, M. P., Singer, D., & Howa, H. (2022). The marine influence index (MII): a tool to assess estuarine intertidal mudflat environments for the purpose of foraminiferal biomonitoring. Water, 14(4), 676. 10.3390/w14040676

60. Kaminski, M. A., Amao, A., Babalola, L., Khamsin, A. B., Fiorini, F., Garrison, A. M., Gull, H. M., Johnson, R. L., Tawabini, B., & Frontalini, F. (2021). Substrate temperature as a primary control on meiofaunal populations in the intertidal zone: A dead zone attributed to elevated summer temperatures in eastern Bahrain. Regional Studies in Marine Science, 42, 101611. 10.1016/j.rsma.2021.101611

61. Kemp, A. C., Buzas, M. A., Horton, B. P., & Culver, S. J. (2011). Influence of patchiness on modern salt-marsh foraminifera used in sea-level studies (North Carolina, USA). The Journal of Foraminiferal Research, 41(2), 114–123. 10.2113/gsjfr.41.2.114

62. Kemp, A. C., Horton, B. P., & Culver, S. J. (2009a). Distribution of modern salt-marsh foraminifera in the Albemarle–Pamlico estuarine system of North Carolina, USA: Implications for sea-level research. Marine Micropaleontology, 72(3-4), 222–238. 10.1016/j.marmicro.2009.06.002

63. Kemp, A. C., Horton, B. P., Vann, D. R., Engelhart, S. E., Pre, C. A. G., Vane, C. H., Nikitina, D., & Anisfeld, S. C. (2012). Quantitative vertical zonation of salt-marsh foraminifera for reconstructing former sea level; an example from New Jersey, USA. Quaternary Science Reviews, 54, 26–39. 10.1016/j.quascirev.2011.09.014

64. Kemp, A. C., & Telford, R. J. (2015). Transfer functions. In Handbook of sea-level research (pp. 470–499). Wiley.

65. Kemp, A. C., Telford, R. J., Horton, B. P., Anisfeld, S. C., & Sommerfield, C. K. (2013). Reconstructing Holocene sea level using salt-marsh foraminifera and transfer functions: lessons from New Jersey, USA. Journal of Quaternary Science, 28(6), 617–629. 10.1002/jqs.2657

66. Khan, N. S., Ashe, E., Horton, B. P., Dutton, A., Kopp, R. E., Brocard, G., Engelhart, S. E., Hill, D. F., Peltier, W., & Vane, C. H. (2017). Drivers of Holocene sea-level change in the Caribbean. Quaternary Science Reviews, 155, 13–36. 10.1016/j.quascirev.2016.08.032

67. Koukousioura, O., Dimiza, M. D., Triantaphyllou, M. V., & Hallock, P. (2011). Living benthic foraminifera as an environmental proxy in coastal ecosystems: A case study from the Aegean Sea (Greece, NE Mediterranean). Journal of Marine Systems, 88(4), 489–501. 10.1016/j.jmarsys.2011.06.004

68. Lance, R. F., Klymus, K. E., Richter, C. A., Farrington, H. L., Carr, M. R., Thompson, N., Chapman, D. C., & Baerwaldt, K. L. (2017). Experimental observations on the decay of environmental DNA from bighead and silver carps. Management of Biological Invasions, 8(3). 10.3391/mbi.2017.8.3.08

69. Langlet, D., Bouchet, V. M., Riso, R., Matsui, Y., Suga, H., Fujiwara, Y., & Nomaki, H. (2020). Foraminiferal ecology and role in nitrogen benthic cycle in the hypoxic southeastern Bering Sea. Frontiers in Marine Science, 7, 582818. 10.3389/fmars.2020.582818

70. Le Cadre, V. R., Debenay, J.-P., & Lesourd, M. (2003). Low pH effects on Ammonia beccarii test deformation: implications for using test deformations as a pollution indicator. The Journal of Foraminiferal Research, 33(1), 1–9. 10.2113/0330001

71. Lee, S. (2000). Carbon dynamics of Deep Bay, eastern Pearl River estuary, China. II: Trophic relationship based on carbon-and nitrogen-stable isotopes. Marine Ecology Progress Series, 205, 1–10. 10.3354/meps205001

72. Legendre, P., & Gallagher, E. D. (2001). Ecologically meaningful transformations for ordination of species data. Oecologia, 129, 271–280. 10.1007/S004420100716

73. Lei, Y., Li, T., Jian, Z., & Nigam, R. (2017). Taxonomy and distribution of benthic foraminifera in an intertidal zone of the Yellow Sea, PR China: Correlations with sediment temperature and salinity. Marine Micropaleontology, 133, 1–20. 10.1016/j.marmicro.2017.04.005

74. Lejzerowicz, F., Esling, P., Majewski, W., Szczuciński, W., Decelle, J., Obadia, C., Arbizu, P. M., & Pawlowski, J. (2013a). Ancient DNA complements microfossil record in deep-sea subsurface sediments. Biology letters, 9(4), 20130283. 10.1098/rsbl.2013.0283

75. Lejzerowicz, F., Esling, P., & Pawlowski, J. (2014). Patchiness of deep-sea benthic Foraminifera across the Southern Ocean: Insights from high-throughput DNA sequencing. Deep-Sea Research Part Ii-Topical Studies in Oceanography, 108, 17–26. 10.1016/j.dsr2.2014.07.018

76. Li, M., Cao, H., Hong, Y.-G., & Gu, J.-D. (2011). Seasonal Dynamics of Anammox Bacteria in Estuarial Sediment of the Mai Po Nature Reserve Revealed by Analyzing the 16S rRNA and Hydrazine Oxidoreductase (hzo) Genes. Microbes and environments, 26(1), 15–22. 10.1264/jsme2.ME10131

77. Li, Q., Lei, Y., Li, H., & Li, T. (2023). Distinct responses of abundant and rare foraminifera to environmental variables in the Antarctic region revealed by DNA metabarcoding. Frontiers in Marine Science, 10, 1089482. 10.3389/fmars.2023.1089482

78. Li, Q., Wong, F. K. K., & Fung, T. (2019). Classification of mangrove species using combined WordView-3 and LiDAR data in Mai Po nature reserve, Hong Kong. Remote Sensing, 11(18), 2114. 10.3390/rs11182114

79. Liu, Z., Khan, N. S., Howard, K. Y. Y., Chung, A., Schweizer, M., & Schunter, C. (2025). Foraminiferal environmental DNA reveals late Holocene sea-level changes. bioRxiv(669806). 10.1101/2025.08.12.669806

80. Lombard, F., Labeyrie, L., Michel, E., Spero, H. J., & Lea, D. W. (2009). Modelling the temperature dependent growth rates of planktic foraminifera. Marine Micropaleontology, 70(1-2), 1–7. 10.1016/j.marmicro.2008.09.004

81. Luke, S. G. (2017). Evaluating significance in linear mixed-effects models in R. Behavior research methods, 49(4), 1494–1502. 10.3758/s13428-016-0809-y

82. Martin, M. (2011). Cutadapt removes adapter sequences from high-throughput sequencing reads. EMBnet. journal, 17(1), 10–12. 10.14806/ej.17.1.200

83. Matsuoka, S., Sugiyama, Y., Shimono, Y., Ushio, M., & Doi, H. (2021). Evaluation of seasonal dynamics of fungal DNA assemblages in a flow-regulated stream in a restored forest using eDNA metabarcoding. Environmental Microbiology, 23(8), 4797–4806. 10.1111/1462-2920.15669

84. Mazurkiewicz, M., Pawłowska, J., Angeles, I. B., Grzelak, K., Deja, K., Zaborska, A., Pawłowski, J., & Włodarska-Kowalczuk, M. (2024). Sediment DNA metabarcoding and morphology provide complementary insight into macrofauna and meiobenthos response to environmental gradients in an Arctic glacial fjord. Marine Environmental Research, 198, 106552. 10.1016/j.marenvres.2024.106552

85. Murray, J. W. (1992). Ecology and distribution of benthic foraminifera: a review. In Y. Takayanagi, and T. Saito (eds.), Studies in benthic foraminifera (Vol. 90, pp. 33–41). Japan: Tokai University Press.

86. Murray, J. W., & Alve, E. (2000). Major aspects of foraminiferal variability (standing crop and biomass) on a monthly scale in an intertidal zone. The Journal of Foraminiferal Research, 30(3), 177–191. 10.2113/0300177

87. Murray, J. W., & Bowser, S. S. (2000). Mortality, protoplasm decay rate, and reliability of staining techniques to recognize ‘living’foraminifera: a review. The Journal of Foraminiferal Research, 30(1), 66–70. 10.2113/0300066

88. Nagler, M., Podmirseg, S. M., Ascher-Jenull J., Sint, D., & Traugott, M. (2022). Why eDNA fractions need consideration in biomonitoring. Molecular Ecology Resources, 22(7), 2458–2470. 10.1111/1755-0998.13658

89. Nevers, M. B., Przybyla-Kelly, K., Shively, D., Morris, C. C., Dickey, J., & Byappanahalli, M. N. (2020). Influence of sediment and stream transport on detecting a source of environmental DNA. Plos One, 15(12), e0244086. 10.1371/journal.pone.0244086

90. Nichols, R. V., Vollmers, C., Newsom, L. A., Wang, Y., Heintzman, P. D., Leighton, M., Green, R. E., & Shapiro, B. (2018). Minimizing polymerase biases in metabarcoding. Molecular Ecology Resources, 18(5), 927–939. 10.1111/1755-0998.12895

91. Oksanen, J., Kindt, R., Legendre, P., O’Hara, B., Stevens, M. H. H., Oksanen, M. J., & Suggests, M. (2007). The vegan package. Community ecology package, 10(631-637), 719. 10.1007/s10531-005-0311-9

92. Pawłowska, J., Lejzerowicz, F., Esling, P., Szczuciński, W., Zajączkowski, M., & Pawlowski, J. (2014). Ancient DNA sheds new light on the Svalbard foraminiferal fossil record of the last millennium. Geobiology, 12(4), 277–288. 10.1111/gbi.12087

93. Pawlowski, J., Esling, P., Lejzerowicz, F., Cedhagen, T., & Wilding, T. A. (2014). Environmental monitoring through protist next-generation sequencing metabarcoding: assessing the impact of fish farming on benthic foraminifera communities. Molecular Ecology Resources, 14(6), 1129–1140. 10.1111/1755-0998.12261

94. Pawlowski, J., Esling, P., Lejzerowicz, F., Cordier, T., Visco, J. A., Martins, C. I., Kvalvik, A., Staven, K., & Cedhagen, T. (2016). Benthic monitoring of salmon farms in Norway using foraminiferal metabarcoding. Aquaculture Environment Interactions, 8, 371–386. 10.3354/aei00182

95. Peres-Neto, P. R., Legendre, P., Dray, S., & Borcard, D. (2006). Variation partitioning of species data matrices: estimation and comparison of fractions. Ecology, 87(10), 2614–2625. 10.1890/0012-9658(2006)87[2614:VPOSDM]2.0.CO;2

96. Piña-Ochoa, E., Høgslund, S., Geslin, E., Cedhagen, T., Revsbech, N. P., Nielsen, L. P., Schweizer, M., Jorissen, F., Rysgaard, S., & Risgaard-Petersen, N. (2010). Widespread occurrence of nitrate storage and denitrification among Foraminifera and Gromiida. Proceedings of the national academy of sciences, 107(3), 1148–1153. 10.1073/pnas.0908440107

97. Pont, D., Rocle, M., Valentini, A., Civade, R., Jean, P., Maire, A., Roset, N., Schabuss, M., Zornig, H., & Dejean, T. (2018). Environmental DNA reveals quantitative patterns of fish biodiversity in large rivers despite its downstream transportation. Scientific Reports, 8(1), 10361. 10.1038/s41598-018-28424-8

98. Prazeres, M., & Pandolfi, J. M. (2016). Effects of elevated temperature on the shell density of the large benthic Foraminifera Amphistegina lobifera. Journal of Eukaryotic Microbiology, 63(6), 786–793. 10.1111/jeu.12325

99. Richirt, J., Riedel, B., Mouret, A., Schweizer, M., Langlet, D., Seitaj, D., Meysman, F. J., Slomp, C. P., & Jorissen, F. J. (2020). Foraminiferal community response to seasonal anoxia in Lake Grevelingen (the Netherlands). Biogeosciences, 17(6), 1415–1435. 10.5194/bg-17-1415-2020

100. Risgaard-Petersen, N., Langezaal, A. M., Ingvardsen, S., Schmid, M. C., Jetten, M. S., Op den Camp, H. J., Derksen, J. W., Pina-Ochoa, E., Eriksson, S. P., & Peter Nielsen, L. (2006). Evidence for complete denitrification in a benthic foraminifer. Nature, 443(7107), 93–96. 10.1038/nature05070

101. Rognes, T., Flouri, T., Nichols, B., Quince, C., & Mahé, F. (2016). VSEARCH: a versatile open source tool for metagenomics. PeerJ, 4, e2584. 10.7717/peerj.2584

102. Romano, E., Bergamin, L., Di Bella, L., Frezza, V., Pierfranceschi, G., Marassich, A., & Provenzani, C. (2021). Benthic foraminifera as environmental indicators in extreme environments: The marine cave of Bue Marino (Sardinia, Italy). Ecological Indicators, 120, 106977. 10.1016/j.ecolind.2020.106977

103. Rush, G., McDarby, P., Edwards, R., Milker, Y., Garrett, E., & Gehrels, W. R. (2021). Development of an intertidal foraminifera training set for the North Sea and an assessment of its application for Holocene sea-level reconstructions. Marine Micropaleontology, 169, 102055. 10.1016/j.marmicro.2021.102055

104. Sawai, Y., Horton, B. P., Kemp, A. C., Hawkes, A. D., Nagumo, T., & Nelson, A. R. (2016). Relationships between diatoms and tidal environments in Oregon and Washington, USA. Diatom research, 31(1), 17–38. 10.1080/0269249X.2015.1126359

105. Schönfeld, J., Alve, E., Geslin, E., Jorissen, F., Korsun, S., & Spezzaferri, S. (2012). The FOBIMO (FOraminiferal BIo-MOnitoring) initiative—Towards a standardised protocol for soft-bottom benthic foraminiferal monitoring studies. Marine Micropaleontology, 94, 1–13. 10.1016/j.marmicro.2012.06.001

106. Scott, D., & Medioli, F. (1978). Vertical zonations of marsh foraminifera as accurate indicators of former sea-levels. Nature, 272(5653), 528–531. 10.1038/272528a0

107. Scott, D. B., Medioli, F. S., & Schafer, C. T. (2007). Monitoring in coastal environments using foraminifera and thecamoebian indicators. Cambridge University Press.

108. Singer, D., Fouet, M. P., Schweizer, M., Mouret, A., Quinchard, S., & Jorissen, F. J. (2023). Unlocking foraminiferal genetic diversity on estuarine mudflats with eDNA metabarcoding. Science of the Total Environment, 902, 165983. 10.1016/j.scitotenv.2023.165983

109. Smith, K. E., Terrano, J. F., Khan, N. S., Smith, C. G., & Pitchford, J. L. (2021). Lateral shoreline erosion and shore-proximal sediment deposition on a coastal marsh from seasonal, storm and decadal measurements. Geomorphology, 389, 107829. 10.1016/j.geomorph.2021.107829

110. Spear, S. F., Groves, J. D., Williams, L. A., & Waits, L. P. (2015). Using environmental DNA methods to improve detectability in a hellbender (Cryptobranchus alleganiensis) monitoring program. Biological conservation, 183, 38–45. 10.1016/j.biocon.2014.11.016

111. Strickler, K. M., Fremier, A. K., & Goldberg, C. S. (2015). Quantifying effects of UV-B, temperature, and pH on eDNA degradation in aquatic microcosms. Biological conservation, 183, 85–92. 10.1016/j.biocon.2014.11.038

112. Sun, Y., Xing, J., Zong, Y., & Hong, G. (2017). Coastal responses and shoreline restoration after Holocene sea-level changes in Hong Kong, China. In Hydraulic Engineering V (pp. 115–120). CRC Press.

113. Tan, F., Khan, N. S., Li, T., Meltzner, A. J., Majewski, J., Chan, N., Chutcharavan, P. M., Cahill, N., Vacchi, M., & Peng, D. (2023). Holocene relative sea-level histories of far-field islands in the mid-Pacific. Quaternary Science Reviews, 107995. 10.1016/j.quascirev.2023.107995

114. Torti, A., Jørgensen, B. B., & Lever, M. A. (2018). Preservation of microbial DNA in marine sediments: insights from extracellular DNA pools. Environmental Microbiology, 20(12), 4526–4542. 10.1111/1462-2920.14401

115. Troth, C. R., Sweet, M. J., Nightingale, J., & Burian, A. (2021). Seasonality, DNA degradation and spatial heterogeneity as drivers of eDNA detection dynamics. Science of the Total Environment, 768, 144466. 10.1016/j.scitotenv.2020.144466

116. Urabe, H., Mizumoto, H., Tsuda-Yamaguchi, F., & Araki, H. (2025). Spatial heterogeneity of eDNA concentration as a predictor of small biomass of fish in a mountain stream. Limnology, 26(1), 223–233. 10.1007/s10201-024-00772-7

117. Walker, J. S., Cahill, N., Khan, N. S., Shaw, T. A., Barber, D., Miller, K. G., Kopp, R. E., & Horton, B. P. (2020). Incorporating temporal and spatial variability of salt-marsh foraminifera into sea-level reconstructions. Marine Geology, 429, 106293. 10.1016/j.margeo.2020.106293

118. Walton, W. R. (1952). Techniques for recognition of living foraminifera. Contribution of Cushman Foundation Foraminiferal Research, 3(2), 56–60.

119. Wang, Y.-F., Feng, Y.-Y., Ma, X., & Gu, J.-D. (2013). Seasonal dynamics of ammonia/ammonium-oxidizing prokaryotes in oxic and anoxic wetland sediments of subtropical coastal mangrove. Applied microbiology and biotechnology, 97(17), 7919–7934. 10.1007/s00253-012-4510-5

120. Welch, W. J. (1990). Construction of permutation tests. Journal of the American Statistical Association, 85(411), 693–698. 10.1080/01621459.1990.10474929

121. Wood, S. A., Biessy, L., Latchford, J. L., Zaiko, A., von Ammon, U., Audrezet, F., Cristescu, M. E., & Pochon, X. (2020). Release and degradation of environmental DNA and RNA in a marine system. Science of the Total Environment, 704, 135314. 10.1016/j.scitotenv.2019.135314

122. Woodroffe, S. A., Horton, B. P., Larcombe, P., & Whittaker, J. E. (2005). Intertidal mangrove foraminifera from the central Great Barrier Reef shelf, Australia: implications for sea-level reconstruction. Journal of Foraminiferal Research, 35(3), 259–270. 10.2113/35.3.259

123. WWF. (2020). Mai Po Nature Reserve and the Ramsar Convention. WWF Hong Kong.

124. Xiong, H., Zong, Y., Qian, P., Huang, G., & Fu, S. (2018). Holocene sea-level history of the northern coast of South China Sea. Quaternary Science Reviews, 194, 12–26. 10.1016/j.quascirev.2018.06.022

125. Yang, K., Wang, L., Cao, X., Gu, Z., Zhao, G., Ran, M., Yan, Y., Yan, J., Xu, L., & Gao, C. (2022). The origin, function, distribution, quantification, and research advances of extracellular DNA. International Journal of Molecular Sciences, 23(22), 13690. 10.3390/ijms232213690

126. Youssef, M., El-Sorogy, A., Al-Kahtany, K., & Saleh, M. (2021). Benthic foraminifera as bio-indicators of coastal marine environmental contamination in the Red Sea-Gulf of Aqaba, Saudi Arabia. Bulletin of Environmental Contamination and Toxicology, 106(6), 1033–1043. 10.1007/s00128-021-03192-w

127. Yu, H. K. Y., Khan, N. S., Desianti, N., Garrett, E., Planavsky, N. J., & Ahmed, A. (2025). The utility of mangrove foraminifera, diatoms, and stable carbon isotope and C/N geochemistry in relative sea-level reconstruction in the Pearl River Delta, China. Marine Geology, 107610. 10.1016/j.margeo.2025.107610

128. Yu, S., Luo, Y., Wu, C., & Xu, W. (2023). Subseasonal variations of convective and microphysical characteristics of extreme precipitation over the Pearl River Delta at monsoon coast. Journal of Geophysical Research: Atmospheres, 128(3), e2022JD037804. 10.1029/2022JD037804

129. Yun, K.-W., Son, H.-S., Seong, M.-J., & Kim, M.-C. (2023). Propidium Monoazide based selective iDNA monitoring method improves eDNA monitoring for harmful algal bloom Alexandrium species. Frontiers in Marine Science, 10, 1257343. 10.3389/fmars.2023.1257343

130. Zhang, J., Kobert, K., Flouri, T., & Stamatakis, A. (2014). PEAR: a fast and accurate Illumina Paired-End reAd mergeR. Bioinformatics, 30(5), 614–620. 10.1093/bioinformatics/btt593

