## Supplementary Figure-Temporal and spatial variability of mudflat and mangrove foraminiferal eDNA assemblages and its implication for sea-level recon for "Temporal and spatial variability of mudflat and mangrove foraminiferal eDNA assemblages and its implication for sea-level reconstruction"

**Other supplementary material includes:** Supplementary Tables 1 to 5 (provided in the excel file "*Supplementary Table - Temporal and spatial variability of mudflat and mangrove foraminiferal eDNA assemblages and its implication for sea-level reconstruction*")

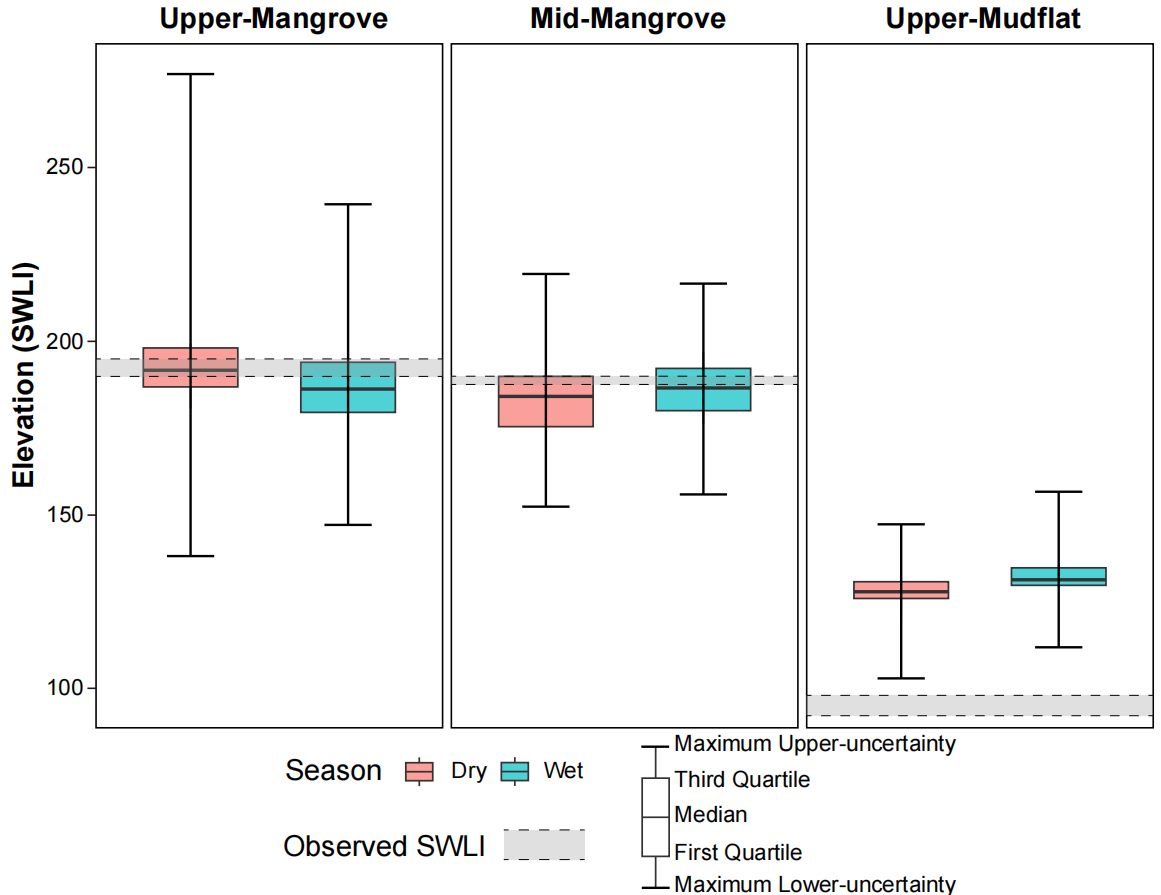

**Figure S1.** Elevation estimates based on foraminiferal eDNA assemblage with Family Globigerinitidae removed. Elevation estimates (Standardized Water Level Index, SWLI) for each monitoring station during dry and wet seasons. Distributions represent six replicate samples per station and season. Error bars indicate the maximum 2σ uncertainty across replicates. The range of observed elevation at each station is shown by the grey dash bar.
